# Neuropeptidergic regulation of Compulsive Ethanol Seeking in *C. elegans*

**DOI:** 10.1101/2021.10.03.462755

**Authors:** Chinnu Salim, Ann Ke Kan, E. Clare Patterson, Changhoon Jee

**Affiliations:** Dept. of Pharmacology, Addiction Science and Toxicology, University of Tennessee Health Science Center, 71 S Manassas St., Memphis, TN

**Author notes:** To whom correspondence should be addressed. Changhoon Jee, Ph.D, Dept. of Pharmacology, Addiction Science and Toxicology, College of Medicine, UTHSC, 71 S. Manassas St., Suite 217, Memphis, TN, 38103.

**Keywords:** Ethanol, Preference, Compulsion, SEB-3, TKR-1, *C. elegans*, CRF_1_ receptor, Neurokinin receptor

## Abstract

An improved understanding of the molecular basis of alcohol seeking despite the catastrophic consequences of alcohol abuse is likely to enrich our treatments for Alcohol Use Disorders (AUD) and comorbidities. The compulsive seeking is characterized by an imbalance between the superior drive to substance and disruption in control of substance use. To model the development of compulsive engagement of alcohol seeking, we exploit two distinct behavioral programs of *C. elegans* in conflict, ethanol preference and avoidance of aversive stimulus, simultaneously. We demonstrate that *C. elegans* exhibited the recapitulation of the pivotal features of compulsive alcohol seeking in mammals, which are repeated attempts, endurance, and finally aversion-resistant ethanol seeking. We find that the neuropeptide signaling via SEB-3, CRF receptor-like GPCR, facilitates the development of ethanol preference and compels animals to seek ethanol compulsively. Furthermore, our functional genomic approach and behavioral elucidation suggest the interaction between neuropeptidergic signaling, SEB-3 and TKR-1, Neurokinin receptor orthologue, to progress compulsive ethanol seeking behavior.

## Introduction

Alcohol Use Disorder (AUD) is a chronic neurobehavioral disorder. A chronic exposure leads to the development of tolerance contributing to increased consumption (^1^, ^2^, ^3^). Subsequently, during the withdrawal or abstinence, alcohol craving and seeking are reinstated (^4^, ^5^, ^6^). Since these three phases model has provided a inferred understanding of the development of AUD (^7^), development of preference and compulsive seeking, known to be involved in progress of alcohol dependence, have been identified as a pivotal and defining characteristic of AUD and other substance use disorders (^8^, ^9^, ^10^). Furthermore, recent Human genome-wide association study (GWAS) demonstrate that heavy drinking and increased consumption are not sufficient causes of AUD, although binge drinking and increased consumption are key risk factors for AUD (^11^). Hence, the neural substrates underlying compulsive seeking despite catastrophic consequences are crucial components of AUD and comorbidities, but the molecular mechanism remains largely elusive. The compulsive seeking in AUD is characterized by an imbalance between superior drive to alcohol and disruption in control of alcohol use (^12^, ^13^). In order to model this highly complex neuromodulation, we addressed sophisticated behavioral paradigms in the simplest and most completely defined connectome with the advantage of the straightforward genetic, behavioral, and neurophysiological investigation, *C. elegans* (^14^, ^15^, ^16^). In a comparative proteomics study, 83% of the worm proteome was found to have human homologous genes and recent meta-analysis of orthology-prediction methods, about 52.6% of human protein-coding genome has recognizable worm orthologues (^17^, ^18^), allow the worm a suitable model organism for functional validation of human genes.

*C. elegans* also has been shown to be a powerful and a deployable genetic tool to study AUD (^19^, ^20^, ^21^, ^22^, ^23^, ^24^, ^25^). Worms showed comparable physiological effects in similar human blood alcohol levels and develop acute functional tolerance in the presence of ethanol. Remarkably, worms also develop ethanol withdrawal symptom of tremor, which is reduced and abolished by replenishment of ethanol (^22^). Previously, we have demonstrated and characterized a form of ethanol-preference behavior. Ethanol preference is elicited by prolonged exposure to ethanol in *C. elegans*, like mammals (^22^). In this study, we further characterized the ethanol preference and demonstrated that *C. elegans* progresses compulsive ethanol seeking behavior. To evaluate the compulsive seeking, the repetitive/obsessive aspects and aversion-resistant seeking scale were applied, which has been used in human genetic studies for reliable assessing alcohol craving and dependence (^6^, ^26^, ^27^). Subsequently, we discovered that *seb-3* (Secretin family Class B GPCR), a corticotropin-releasing factor (CRF) receptor-like GPCR in *C. elegans*, facilitates the development of ethanol preference and progress to compulsive ethanol seeking behavior. A CRF receptor has long been known to have roles in brain stress response and been implicated in the pathophysiology of anxiety and AUD (^28^, ^29^, ^30^, ^31^). Additionally, its function has been implicated in compulsive drug self-administration and stress-induced reinstatement of drug seeking in mammalian studies (^32^). Furthermore, a recent GWAS identified CRFR1 loci is associated with habitual alcohol intake (^33^). We have analyzed differentially expressed genes in a gain-of-function variant of *seb-3*, that facilitates faster development of preference and enhances compulsive seeking of ethanol. Here, we suggest TKR-1, neurokinin receptor orthologue, is upregulated by potentiation of SEB-3 to progress maladaptive compulsive ethanol seeking affecting susceptibility to AUD.

## Materials and Methods

All strains were maintained on nematode growth media (NGM) plates with Escherichia coli (OP50) at 20°C (^34^) and the hermaphrodite was used for behavioral analysis. The wild-type animals used for the experiment were the Bristol N2 strain. The strains *tkr-1*(*ok2886*), *tkr-2* (*ok1620*) and *tkr-3(ok381)* were obtained from Caenorhabditis Genetics Center (CGC, Minneapolis, MN, USA), which is supported by the National Institutes of Health - Office of Research Infrastructure Programs (P40 OD010440). *seb-3*(*eg696*)*gf* previously isolated from a genetic screening and the *seb-3(tm1848)lf* strain was obtained from S. Mitani (Tokyo Women’s Medical University, Tokyo, Japan) and backcrossed twice with N2 (^22^).

### Trajectory Analysis of *C*.*elegans* locomotion in the development of ethanol preference

NGM was added to each well of 4 well tissue culture dish (1.9 cm2) to fill all the wells equally to the top. Outside of well also filled up with NGM and the boundary were slightly covered by disruption of surface tension when NGM was not solidified, which contained a gradient of ethanol but an allowance of free moving of worms on the surface. Out of the 4 wells, one well was selected and the glass Pasteur pipette (2ml volume) punctured to make a hole for adding ethanol up to 300Mm. The plate was sealed with parafilm after adding ethanol then 1 hour later used. 1day adult animals were washed twice in S-basal (100mM NaCl, 50mM potassium phosphate (pH 6.0), ^35^) and once in distilled water. After a final wash, 10 worms (Naïve or Ethanol treated) were introduced to the mid-region of 4-well plate by gentle picking with a platinum wire. Seal with parafilm and recorded the locomotion for 30 minutes in Wormlab (MBF Bioscience). The trajectory and time spent in the distinct area were analyzed by Wormlab software (MBF Bioscience). For the control speed and travel distance, 10 Naïve or Ethanol treated animals were placed on NGM plate with OP50 lawn (food) then 15 min locomotion was recorded and analyzed with Wormlab (MBF Bioscience).

### Aversion-resistant Ethanol seeking Assay

The ethanol pre-exposure plates were prepared as previously described (^19^, ^21^). Preexposure plates were 6-cm NGM plates that had been seeded with bacteria on half of the plate and were dried for 2 h. Ice-cold ethanol was added up to final concentration of 300 mM ethanol (from our previous publication; ^21^) and worms were introduced to ethanol plates for 4 hours pre-exposure with parafilm sealing. Briefly, well-fed naïve or ethanol pretreated animals (300mM) were washed twice with S-buffer [100mM NaCl, 50mM potassium phosphate (pH 6.0)] and once in distilled water. Then animals were placed on mark above B (arrow in fig. 2) at the chemotaxis assay plate (^36^,^37^). Animals were allowed to move freely on an assay plate with different concentrations of CuSO_4_ barrier to see the compulsive alcohol seeking behavior. The concentration of copper barrier was taken from the previous publication of Hilliard et al, 2004 (^38^), starting from a lower concentration of 2 mM to 20 mM. Before making the chemical barrier, allow the chemotaxis plates to dry for 1 hr. To make a chemical barrier, pre-cut (3mm thickness) Whatman filter paper (cat No-3030-335) was placed at the center of the 100 mm chemotaxis assay plate dividing the plate into two equal halves across the midline, as marked aversive barrier in (fig. 2). A 100 µl of CuSO_4_ (each concentration; 2 to 20mm) was added to this filter paper barrier as per the experiment, from one edge to the other end. Briefly after adding CuSO4 solution, the filter paper was gently removed using forceps. Meanwhile, 1 µl of 1M NaN3 was added to the point marked A (fig. 2) and 30 µl of ethanol (200 proof) on top of it. Immediately after the chemical compound was absorbed into the 10cm chemotaxis plate, about 100 to 150 washed animals were placed in the area marked in B section using glassware micropipette. The excess liquid was removed with a Kimwipe for animals not to be clumped at the origin. 40 minutes later with parafilm sealing, the number of animals in each section marked (Fig.2; A, B, C and D) was counted to calculate the Seeking index. The index was calculated by [(number of animals in A - number of animals in B)/Total number of animals [Seeking index SI= (A-B)/Total(A+B+C+D)]. To measure the copper sensitivity, 20 naïve or Ethanol-pretreated animals were placed on the aversion-resistant assay plate with 5mM CuSO_4_ barrier then the locomotion was recorded for 30min. Time to respond to the copper barrier, which was stopping forward locomotion and backward then turn to avoid it, was measured in each animal’s first encounter with the copper barrier. To assess with different types of aversive stimuli than copper, 5mM or 10mM Denatonium benzoate was applied to the assay plate in the same way to create an aversive barrier. SI in each trial was obtained from the population assay of 100 to 150 animals. Avoidance assay with drop test (0.1mM, 1mM, and 5mM CuSO_4_) was conducted as shown in previous study (^39^, ^38^), to measure the sensitivity to nociceptive stimuli of WT and *seb-3(eg696)* gf animals. Single L4 worms were transferred to each NGM plate with abundant OP50, and 15 h later the assay was conducted. Assays were conducted under blind conditions and were not performed more than once on any individual animal. The avoidance index (AI) in (b) is the number of positive responses divided by the total number of trials. Mean latency to respond was calculated for all positive responses.

### Sequence alignment

TKR-1 protein sequences (NP_499064.2) were analyzed by database similarity search (^40^) and the multiple protein sequences were simultaneously aligned using the COBALT, a constraint based alignment tool (^41^). The transmembrane helix domains were predicted using the conserved domain database (CDD) (^42^). For the phylogenetic tree analysis, Worm, human, rat, and mouse neurokinin receptor sequences used for alignment: worm TKR-1 (NP_499064.2), h_TACR1/NK1R (NP_001049.1), h_TACR2/NK2R (NP_001048.2), h_TACR3/NK3R (NP_001050.1), r_TACR1/NK1R (NP_036799.1), r_TACR2/NK2R (NP_542946.1), r_TACR3/NK3R (NP_058749.1), m_TACR1/NK1R (XP_006505926.1), m_TACR2/NK2R (NP_067357.1), and m_TACR3/NK3R (NP_033340.3). The phylogenetic tree was constructed by COBALT using minimum evolution method.

### Statistical Analysis

The Mean and standard error of the mean (SEM) were determined for all experimental parameters. The data were analyzed employing the Chi-square test or Mann-Whitney test (Graph pad prism version 8.0.1). Values below 0.05 were considered as significant.

### Microarray Analysis

Isolation of nucleic acid and microarray followed by analysis of differentially expressed gene profiling was done in Molecular Recourse center of excellence (MRC), UTHSC. Briefly, synchronized 1 day adult population of WT and *seb-3gf(eg696*) where used for total RNA isolation. Approximately 10 one day adult animals where allowed to lay eggs per 15 NGM plates seeded with OP 50 for 8 to 10 hours. After 10 hours, the adult animals were removed and embryo where allowed to grow at 20 degree. After 72 hours, one-day adult animals from each plate were washed out with M9 buffer and washed the worms three times before collected in 1.5ul nuclease free sterile Eppendorf tubes and stored at -80 with RNA later until use. Isolation of nucleic acid (total RNA) was done in Qi cube unit using Qiagen RNeasy mini kit according to manufacturer’s instruction and standard protocol. Quantification of nucleic acid was done by Agilent2100 bioanalyzer, nanodrop and qubit. Individual sample (*n-3*) was run for Affymetrix microarray focusing on individual response. Analysis was done using Affymetrix Genotyping Console and Affymetrix Transcriptome Analysis Console software.

For the population of germ cell production inhibited, 5’-fluorodeoxyuridine (FUdR) was treated from late L4 till the collection for the RNA preparation. The total RNA was isolated from 3 replicates of synchronized 1-day adult population of WT, *seb-3gf(eg696*), FUdR-treated WT, and FUdR-treated *seb-3gf(eg696*), respectively. Since FUdR affects some physiological function such as osmotic stress related life span extension (^43^), gene expression profiling from germ cell production inhibited was verified by excluding overlapping hits through cross-validation through mutual comparison. 1 candidate gene that changed in meaningful direction and amount by FUdR treatment was removed from the final differentially expressed genes candidates. RNA was reverse transcribed and labeled using GeneChip™ Whole Transcript (WT) Pico kit and hybridized to GeneChip™ C. elegans Gene 1.0 ST Array (Thermo Fisher Scientific) for whole-genome expression profiling.

Text files were retrieved from UTHSC Molecular Resource Center after normalization performed by Affymetrix Expression Condole. Quality assurance was checked against reference probes to ensure quality of data. Gene names, accession numbers, and expression were mined from each text file for each sample. All non-annotated information was removed from the file leaving only annotated gene expression. A student *t* test was run for pairwise interactions in order to obtain pvalues for significance. Only genes with a pvalue < 0.05 were considered significant. The mean, variance, standard deviation, and fold change were calculated for each pairwise comparison. Benjamini Hochberg false discovery rate method was applied in order to obtain the adjusted pvalue for each gene (^44^). Heatmaps were created to visualize the expression of the significant genes of each pairwise comparison. Only genes with a p value < 0.05 were considered significant (^44^). Gene ontology enrichment analysis was conducted in the GO resource (GO; http://geneontology.org) and WormBase (https://wormbase.org/tools/enrichment/tea/tea.cgi) and verified through cross-validation through mutual comparison (^45^, ^46^, ^47^, ^48^, ^49^). Additionally, a comprehensive comparison of the results performed in Tissue Enrichment Analysis (TEA), Phenotype Enrichment Analysis (PEA) provided by WormBase was used to investigate genes predicted them to function in the same cellular pathway for further functional evaluation (^48^, ^49^).

## Results

An ethanol preexposure elicits ethanol preference in wild-type (WT) *C. elegans* animals. Previous quadrant-plate assay successfully quantified that Ethanol pretreated animals preferentially accumulated in the ethanol area for 30 minutes of permission to move freely, whereas naïve animals accumulated more in the non-ethanol area at first experience (^21^). Here, we further investigated the comprehensive response of *C. elegans* to the ethanol including initial response. WT animals were allowed to move freely on the assay plate, which contains ethanol in only one of the 4 well (1.9 cm^2^) of the culture dish (Fig.1) and recorded locomotion of animals were analyzed. The ethanol concentration was determined based on the behavioral and physiological response to various concentrations shown in previous studies (^19^, ^20^, ^21^). It has been shown that the internal concentration of ethanol for WT animals, which were exposed and forced to be in 400mM ethanol plate, were comparable with 0.1% blood alcohol in human, corresponds to 21.7 mM ethanol in the blood (^19^, ^20^). We treated and tested animals in 300mM ethanol, which was shown to induce ethanol preference of WT animals within 4 hours while being rapidly sobered from intoxication when treated chronically (^21^). The trajectory, exploring patterns, of an individual animal exhibited a distinct difference of response to ethanol between naïve and pretreated animals (Fig. 1). Naïve animals kept exploring the area with and without ethanol when the animal is introduced to the assay plate which is only one of the four wells contains ethanol. When animals first encountered ethanol and moved in the ethanol area, the speed of locomotion was reduced which was corresponding to the behavioral characteristic of intoxication. Then, they left the ethanol zone and kept moving, exploring in the non-ethanol zone more time (Fig. 1a and Fig. 1c; 0% appearance in the ethanol area of the assay plate for some of the naïve animals such as #4, #5). In contrast, ethanol-pretreated WT animals headed straight to the ethanol zone and stayed in the ethanol zone (Fig. 1b and Fig. 1d). The naïve animals stayed longer in the non-ethanol area (66% of time spent) whereas ethanol pretreated animals spent 89% of the time in the ethanol area (Fig. 1d). First, we addressed the question of whether the change of locomotion behavior is due to an inability to keep moving. Ethanol pretreated (300mM) animals were placed on the non-ethanol NGM and their locomotion was analyzed. The ability to move was not defective after ethanol pretreatment. While animals exposed to 300mM exogenous ethanol for the pretreatment (Fig.1), the speed of locomotion was reduced initially but recovered. Thus, no difficulty was observed in the movement of pretreated animals (Fig. 2a, 2b). Ethanol preexposure produces a reversible change in body bends of worm interlocked with the locomotion resulting in a progressive decrease in the speed of movement (^19^). Subsequently, a decrease of speed is recovered in the presence of ethanol that was shown to be corresponding to intoxication and acute functional tolerance in other systems (^20^). Besides, pretreated animals represented an increase of speed and travel distance upon ethanol withdrawal (Fig. 2c), comparable to behavioral stimulation induced by ethanol withdrawal in other system (^50^).

**Fig.1.**
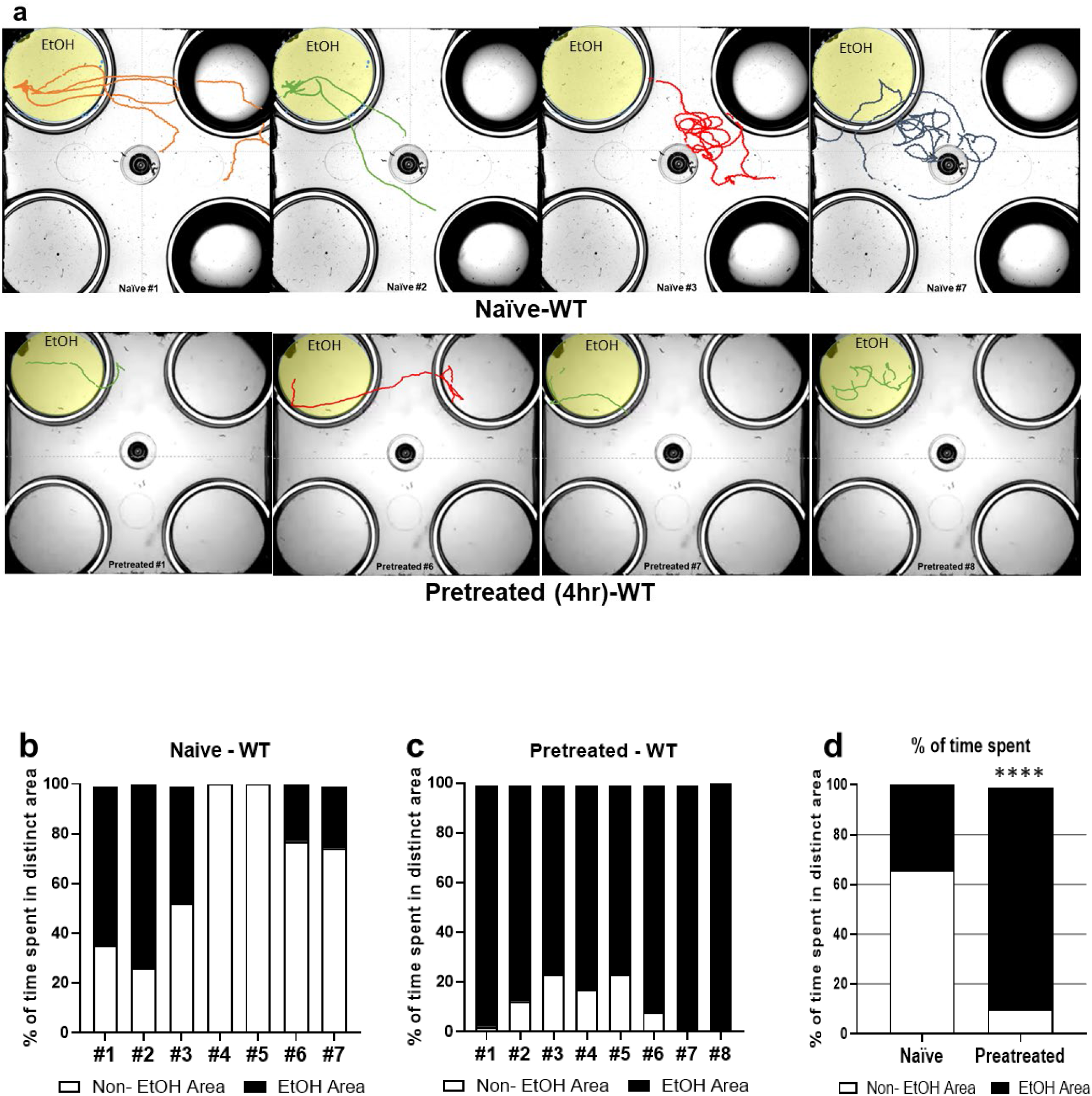
Ethanol-pretreated WT animals headed straight to the ethanol area and remained stayed. (a, b) Trajectory of individual WT animal (Naïve or Ethanol pretreated). The naïve (a) or ethanol pretreated animals (b), respectively, were placed in the middle of assay plate that contains ethanol (300 mM) only in the left top well. All wells are marginally covered by media that allows free moving between the area. Ethanol-pretreated WT animals (4 hours;300mM) headed straight to ethanol area and stayed, whereas naïve wild type animals explored around. (c, e) Behavioral quantification of individual animal (c; Naïve, d; 4hr-Ethanol pretreated) and average of total percentage time spent in the distinct area (e). The ethanol pretreated animals spent more time in ethanol area. The data were analyzed employing Chi-square test. chi-square test indicated; df 60.67, 1 z 7.789, ****P <0.0001

**Fig.2.**
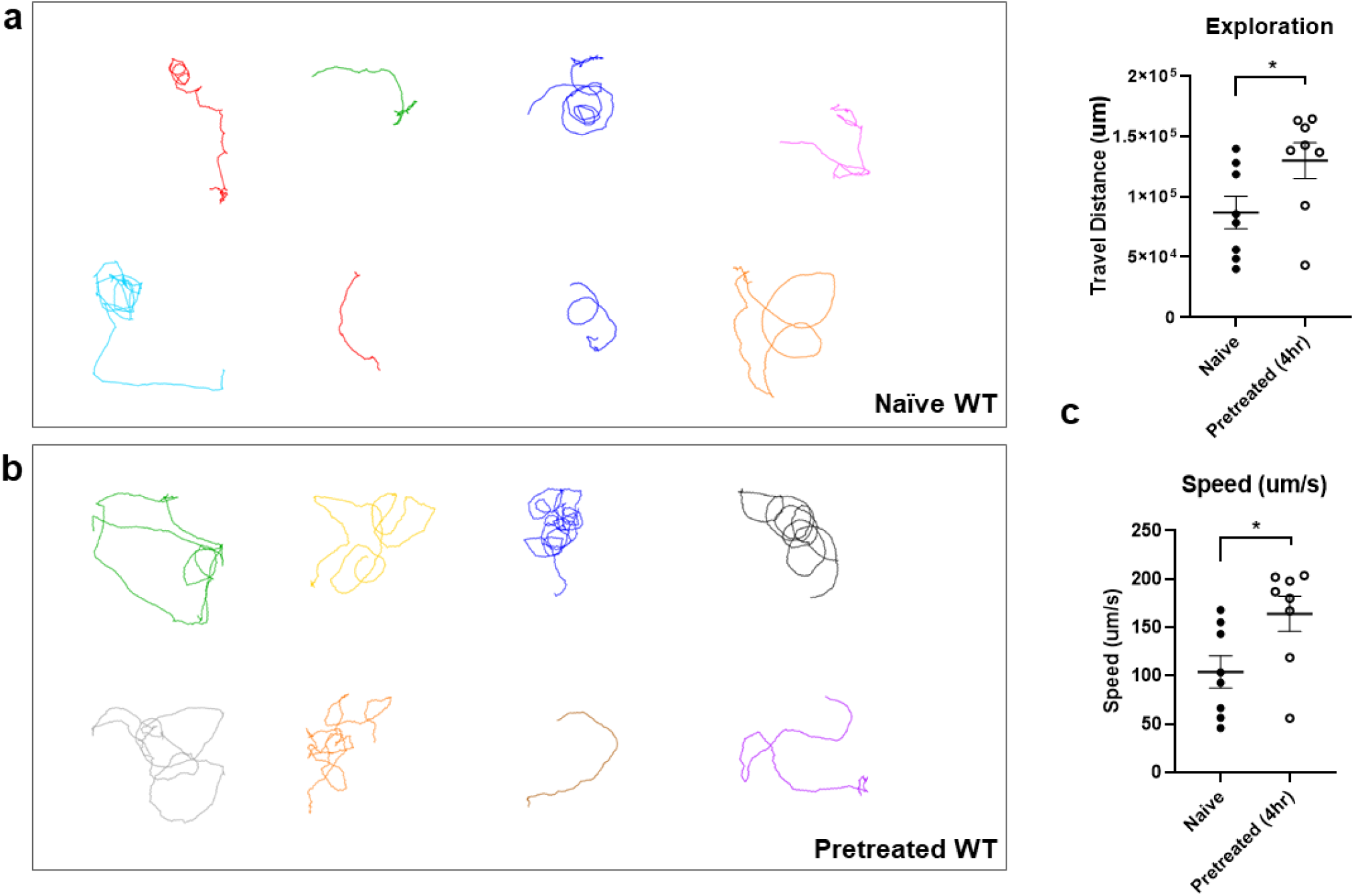
Ethanol pretreated animal is not defective in the locomotion. The locomotion trajectory of naïve (a) or ethanol-pretreated (b) animals is represented. (c) The stimulation of locomotion, increase of speed and travel distance, was observed in ethanol pretreated WT animals.

To further address the question of how preexposure to ethanol might lead animals to stay in the ethanol area although ethanol pretreated animals had an increased exploring and enhanced active behavior, we established the measurement for the motivational strength of ethanol preference progressed by ethanol pretreatment (Fig. 3a). We exploited two distinct behavioral programs in conflict, an ethanol-seeking and avoidance of aversive stimuli blocking ethanol-seeking simultaneously. The motivational strength for the ethanol was indexed by exposing the animals to the ethanol and aversive stimuli at the same time after the pretreatment of ethanol. The nociceptive stimuli of Cu ^2+^ set up as an aversive barrier to interfere with chemotaxis to ethanol (Fig. 3a, 3b, and 3c). *C. elegans* makes use of polymodal sensory neurons to detect a wide range of aversive and painful stimuli including water soluble chemical repellents such as quinine, denatonium benzoate, and Cu ^2+^ (^39^, ^38^) and to let them to avoid it. An animal who detects the aversive stimulus stop its forward locomotion and turn away from the source of stimulus resulting in avoidance. Increased attempt to cross over and endurance of seeking ethanol was observed in the pretreated animals (Fig. 3a). Average trials of single animal (counted out of 103 worms) for 5minutes was 0.56 in naïve whereas 2.28 in pretreated (counted out of 80 worms). These repeated trials of pretreated animals resulted in crossing over the chemical barrier and heading to the ethanol area. The 23.08 % of the trials in pretreated animals resulted in successful crossing to seek ethanol whereas only 3.45 % of the trials in naïve animals. These repetitive attempting and crossing over the aversive barrier were hypothesized to be proportional to motivational compulsion to seek ethanol since ethanol pretreatment does not interfere with the ability of worms to detect Cu ^2+^ barrier. The sensitivity of ethanol-pretreated animals to aversive stimuli was not altered (Fig. 3b). Subsequently, we quantified the animals successfully reached to ethanol over aversive Cu ^2+^ barrier in time and Seeking Index (SI) was obtained as shown in (Fig. 3a). We tested what range of Cu ^2+^ concentration naïve or pretreated WT animals could cross over to reach the ethanol area. The drive of pretreated animals to seek ethanol superseded its aversive Cu ^2+^ response in a concentration-dependent manner (Fig. 3c). The ethanol preference developed by pretreatment enabled WT animals to cross the Cu ^2+^ barrier in low concentration (2mM, 5mM), aversive stimuli that Naïve animal does not cross. However, while going from lower concentration to higher concentration of barrier (10mM, 20mM), ethanol pretreated animals failed to overcome the Cu ^2+^ barrier. 4 hours pretreatment of ethanol was not enough to enable WT animals to cross the aversive Cu ^2+^ barrier to reach the ethanol area. Evidently, this change is strengthened with longer ethanol exposures (data not shown). Then we also introduced another aversive stimulus, denatonium benzoate as a barrier. Without pretreatment of ethanol, all the tested concentrations of denatonium barrier successfully block to interfere the naïve animals’ chemotaxis to ethanol. However, pretreated animals showed endurance against the denatonium barrier to reach the ethanol area and crossed the low concentration of obstacle (Fig. 3d). Since the compulsivity is defined as the urge to perform and persist to take a substance that escapes control despite serious negative consequences (^51^, ^52^, ^53^), these results suggest that aversion-resistant ethanol seeking approach can be used as a proxy measure for the compulsive ethanol seeking behavior.

**Fig.3.**
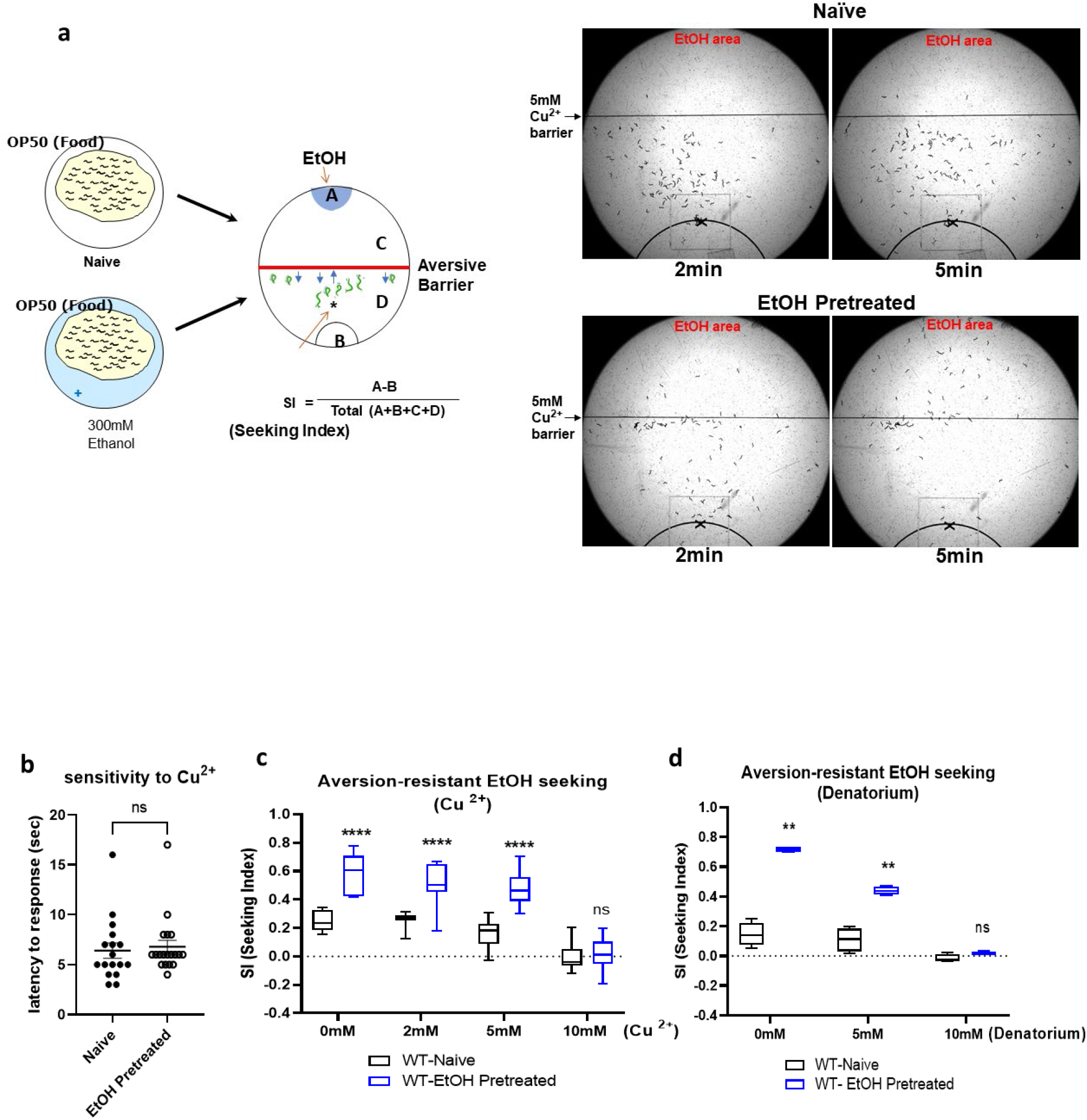
Behavioral quantification of compulsive ethanol seeking after exposure to ethanol against aversive chemical barrier. (a) Diagrammatic representation exhibited experimental design to quantify aversion-resistant ethanol seeking. The different concentrations of Cu^2+^ created aversive barrier without mechanical obstacles to quantify the motivational strength of seeking ethanol behavior in animals. (b) Cu sensitivity of WT animals are not altered after ethanol exposure. (c) EtOH pretreated animals demonstrates more animals cross over the Cu barrier for ethanol (aversion-resistant seeking), in low concentration (2 and 5 mM), than does Naïve animals [F_EtOH pretreated_(1, 18)=46.44, p<0.0001; F_Concentration_(4,72)=99.01, p<0.0001; F _pretreated x Concentration_(4, 72)=11.57, p<0.001]. Moving to higher concentration of copper (10mM), ethanol pretreated WT animals failed to overcome the aversive barrier. A two-way ANOVA comparison of the animal status over concentrations of barrier showed significant differences based on EtOH pretreated, concentrations, and the interaction of the two. Significant post hoc differences (Bonferroni’s test) between naïve and EtOH pretreated animals at no barrier, 2mM, 5mM, and 10mM is shown (p<0.0001, ****). Values are mean ± SEM. N=10 in each conc. (d) EtOH pretreated animals demonstrates more animals cross over the Denatorium barrier for ethanol (aversion-resistant seeking), in low concentration, than does Naïve animals [F_EtOH pretreated_(1, 6)=598.8, p<0.0001; F_Concentration_(1.712, 10.27)=119, p<0.0001; F _pretreated x Concentration_(2, 12)=45.81, p<0.0001]. A two-way ANOVA comparison of the animal status over concentrations of barrier showed significant differences based on EtOH pretreated, concentrations, and the interaction of the two. Significant post hoc differences (Bonferroni’s test) between naïve and EtOH pretreated animals at no barrier, 5mM, and 10mM is shown (p<0.01, **). Values are mean ± SEM. N=4 in each conc.

Neuropeptide signaling orchestrates many complex behaviors in the brain, including compulsive behavior. Previously, we demonstrated the functional conservation of the neuropeptide corticotropin-releasing factor (CRF) system in responses to stress and ethanol in worms and mice (^22^). We reported that *seb-3*, CRF (corticotropin-releasing factor) receptor-like G protein-coupled receptor of *C. elegans* facilitated the development of acute tolerance to ethanol and ethanol withdrawal symptom (^22^). Since the CRF (corticotropin releasing factor) receptor function has been implicated in compulsive drug self-administration and stress-induced reinstatement of drug seeking in mammalian studies, we hypothesized *seb-3(eg696)* gain of function (gf) animals would also be able to exhibit enhanced alcohol seeking against the higher concentration of Cu ^2+^ barrier. A *seb-3(eg696)*, dominant mutation, was suggested and functionally evaluated as a gain of function mutation due to its identity of mutation, which was single amino acid change in a conserved residue at the third intracellular loop, the binding region for G proteins (^22^, ^54^, ^55^, ^56^). First, ethanol preference of *seb-3(eg696)* animals and *seb-3(tm1848)* Knockout animals due to its deletion (^22^) was tested in the preference assay on ethanol-containing quadrant plates (Fig. 4a). Animals were allowed to move freely for 30 minutes between two control quadrants without ethanol and two ethanol quadrants. The naïve animals of WT, *seb-3(eg696), and seb-3(tm1848)* accumulated on non-ethanol regions. No significant ethanol preference was observed in naïve animals of all WT, *seb-3(eg696), and seb-3(tm1848)* (Fig. 4b). Remarkably, *seb-3(eg696)* gf animals represented much faster and greater ethanol preference after ethanol pretreatment. Significant ethanol preference was observed in animals pretreated with ethanol for 2 hours whereas WT animals did not (Fig. 4c). Additionally, greater development of preference was observed in *seb-3(eg696)* animals after 4 hours ethanol pretreatment. Moreover, impaired development of ethanol preference was observed in ethanol pretreated *seb-3(tm1848)* animals (Fig. 4c) suggesting SEB-3 facilitated the ethanol preference. Subsequently, we conducted aversion-resistant ethanol seeking assay with *seb-3* mutant animals. Quantification of *seb3(eg696)* gf mutant animals’ motivational drive to seek ethanol showed enhanced aversion-resistant ethanol seeking behavior compared to WT animals in overall concentrations of barrier including higher concentrations (10mM and 20mM). The ethanol pretreated *seb-3(eg696)* gf animals gathered at the barrier in the middle of assay plate and the repetitive attempting led animals to cross over the aversive barrier. The pretreated *seb-3(eg696)* gf animals exhibited significantly high SI values not only in lower but also in the higher concentration of Cu ^2+^ (Fig. 5a). Nevertheless, sensory perception of aversive Cu ^2+^ stimuli was not defective in *seb-3(eg696)* gf animals (Fig. 5b, 5c, 5d, and 5e) represented enhanced ethanol preference, which could override the interference of noxious stimuli. Although *seb-3(eg696)* gf animals are slightly more sensitive to nociceptive stimuli (^22^, Fig. 5c), crossed the aversive barrier for ethanol. Additionally, *seb-3(tm1848)* Knockout animals did not develop chemotaxis to ethanol after pretreatment as much as WT animals nor cross both low and high concentrations (2mM, 5mM, 10mM) of Cu ^2+^ barrier for ethanol (Fig. 5f). Taken together with ethanol preference, we conclude SEB-3 also contribute to the progress of compulsive ethanol seeking.

**Fig. 4.**
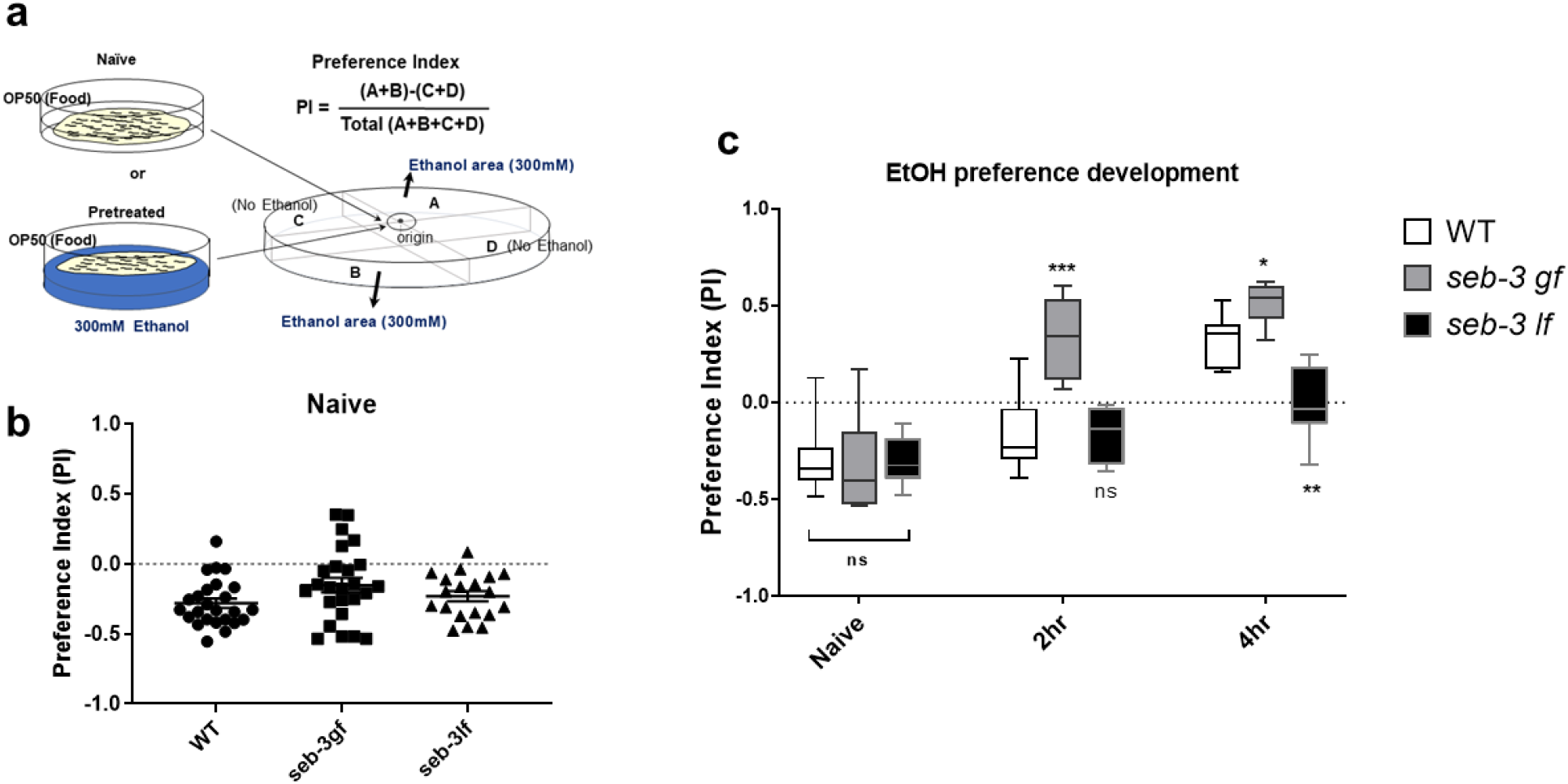
SEB-3 facilitates the development of ethanol preference. (a) Diagrammatic representation of ethanol preference assay. Naïve or ethanol pretreated animals remain free to explore on the quadrants plate before counting. (b) Naïve animals accumulate primarily in the non-ethanol region. One-way ANOVA, P>0.05, F (2, 66) =2.539, ns compared to WT in post hoc multiple comparison test; Dunnett’s. (c) Ethanol preference developed more rapidly and greater in *seb-3gf* animals, whereas impaired in *seb-3lf* animals. [Fgenotype(2, 27)=15.76, p<0.0001; Ftime(1.871, 37.42)=72.56, p<0.0001; FGenotype x time(4, 40)=9.982, p<0.001]. A two-way ANOVA comparison showed significant differences based on genotype, time, and the interaction of the two. Significant post hoc differences (Bonferroni’s multiple comparison test) between the genotypes (WT vs *seb-3gf*) or (WT vs *seb-3if*) at naïve, 2hours, and 4hours are shown (p<0.05, *; p<0.01, **; p<0.0001, ****).

**Fig. 5.**
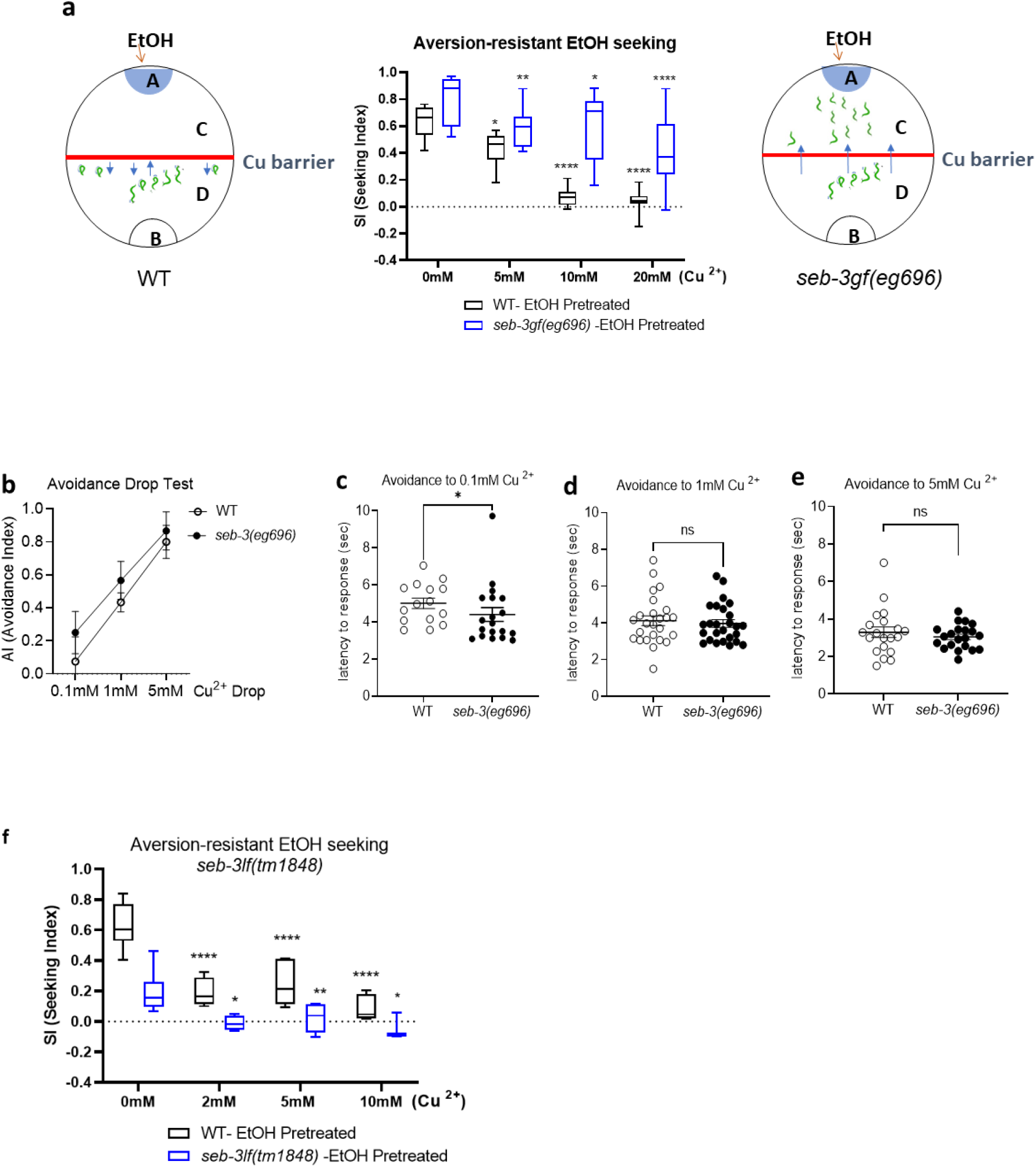
The ethanol pretreated *seb-3gf* animals for 4 hours surmount a stronger aversive barrier to seek ethanol. (a) Strength of ethanol seeking is represented by the SI under different concentration of copper barrier (no barrier, 10mM and 20mM). A *seb-3gf* strain demonstrates more animals cross over the barrier for ethanol (aversion-resistant seeking), in overall concentrations, than does the WT strain [F_Genotype_(1, 18)=35.78, p<0.0001; F_Concentration_(3, 53)=40.32, p<0.0001; F_Genotype x Concentration_(3, 53)=6.586, p<0.001]. A two-way ANOVA comparison of the strains over concentrations of barrier showed significant differences based on genotype, concentrations, and the interaction of the two. Significant post hoc differences (Dunnett’s test) between no barrier versus 5mM, 10mM, or 20mM in each genotype (WT and *seb-3gf* animals) is shown (p<0.05, *; p<0.01, **; p<0.0001, ****). Box and whisker represent minimum to maximum of 10 trials of population assay (N=10). (b-e) Avoidance assay with drop test (0.1mM, 1mM, and 5mM CuSO4). The avoidance index (AI) in (b) is the number of positive responses divided by the total number of trials. The latency to stop forward and initiate backward movement was measured. Data were obtained from 10 or more animals and mean values from 3 or more trials were analyzed by one-way ANOVA with a post-hoc Dunnett’s test; non-significant differences [p=0.0820, F_genotypes_(1, 7)=4.117, p=0.7705; F_concentration x geneotype_(2, 7)=0.2707] (b) or two-tailed t-test; p<0.05, * (c-e). (f) The development of aversion-resistant ethanol seeking is impaired in *seb-3lf* animals. A two-way ANOVA comparison [F_Genotype_(1, 17)=24.64, p<0.0001; F_Concentration_(3, 30)=18.35, p<0.0001; F_Genotype x Concentration_(3, 40)=6.005, p=0.0018]. Significant post hoc differences (Dunnett’s test) between no barrier versus 2mM, 5mM, or 10mM in each genotype (p<0.001, ***; p<0.0001, ****).

Having defined the progress of the compulsive ethanol seeking, we next investigated a gene expression profile of *seb-3(eg696)* gf animals to identity the genes and pathways underlying the enhancement of ethanol preference and progress of compulsive ethanol seeking. The total RNA was isolated from synchronized young adult populations of which germ cell production was pharmacologically inhibited to improve the chance of identification of differentially expressed genes in somatic cells including neuronal tissue, then applied to the Affymetrix gene chip array. Microarray analysis revealed 716 transcripts that are differentially expressed in *seb-3(eg696) gf* animals (≥1.5X, Fig. 6a) compared to WT animals. Subsequent Gene Ontology (GO) enrichment analysis (See Methods) identified total 16 GO terms in upregulated and downregulated gene clusters (Fig. 6b, 6c, Sup.1, and Sup.2). The most prominent enrichment in biological processes includes the immune system process, defense response, and response to biotic stimulus, which is in line with the role of the stress response for survival and reproduction in adverse conditions.

**Fig. 6.**
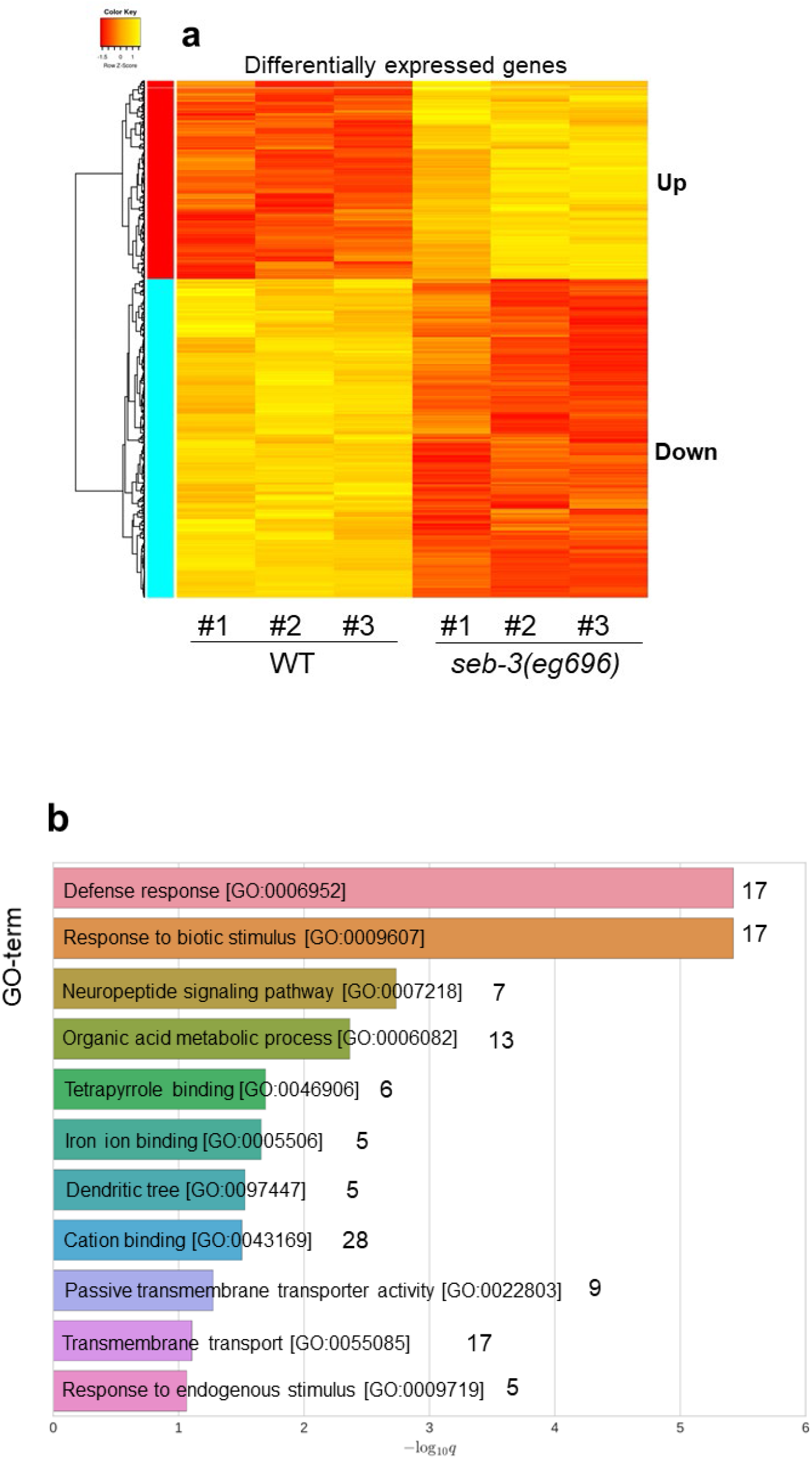
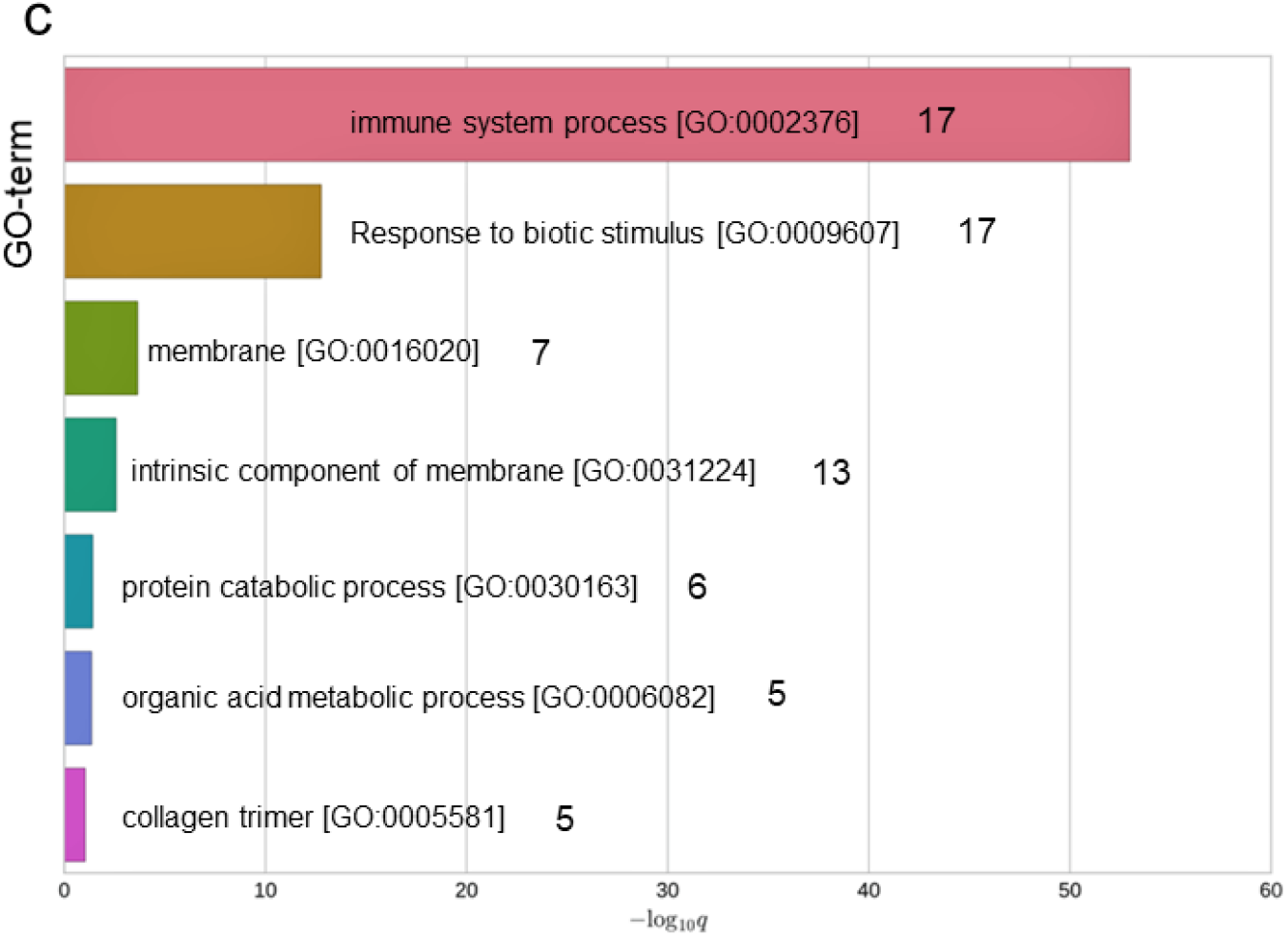
Differentially expressed gene profiling of genetic variant vulnerable to compulsive seeking behavior. (a) Altered expression of transcripts in *seb-3gf*. Gene Ontology (GO) enrichment analysis revealed 11 GO terms that upregulated genes are enriched (b) and 7GO terms of downregulated genes (c).

Remarkably, we identified the upregulated *tkr-1* in Neuropeptide signaling pathway (GO:0007218, Fig. 6b, Sup. 2). A *tkr-1* is predicted to encode a G protein-coupled receptor (GPCR), which is a human orthologue of Tachykinin Receptor also known as neurokinin receptor (^18^). Tachykinins/Neurokinins and their receptors are neuropeptides signaling conserved from invertebrates to mammals (^57^, ^58^). In mammals, tachykinins are widely distributed in the nervous systems and gastrointestinal tract. The three genes encoding precursors of tachykinins give rise to nine distinct tachykinins, which interact with three different receptors such as neurokinin 1 receptor (NK1R), neurokinin 2 receptor (NK2R), and neurokinin 3 receptor (NK3R) (^59^). TKR-1 shows 51% similarity with human NK1R in overall 406 amino acids and 52% similarity with human NK3R. TKR-1 represents the putative ligand-binding pocket defined according to the NMR evidence of Substance P (SP) docking to the NK1R pocket (Fig.7a, ^60^), The crystal structure of NK1R in complex with clinically used antagonist also endorse the important role of this region, that is a ligand bind deeper within the receptor core (^61^). Tachykinin/Neurokinin system is functionally pleiotropic and has been reported to be involved in the stress responses, anxiety, pain, inflammation, immune response, and sensory processing (^59^, ^62^, ^63^, ^64^, ^65^, ^66^, ^67^). For the functional evaluation of *tkr-1* in association with the ethanol preference and compulsive ethanol seeking, we tested the KO mutant of *tkr-1 (ok2886)* in the assay for aversion-resistant ethanol seeking. The *C. elegans* genome includes 3 genes predicted to encode Neurokinin receptor family proteins, *tkr-1, tkr-2*, and *tkr-3* as shown in the phylogenetic analysis of the Tachykinin/Neurokinin Receptor family (Fig. 7b). The phylogenetic tree was constructed by COBALT (constraint-based alignment tool for multiple protein sequences) using minimum evolution method (^25^). We also tested KO mutant animals of *tkr-2 (ok1620)* and *tkr-3(ok381)* in the neurokinin receptor family. The tested are putative KO strains due to the deletion of genomic loci (Fig. 7c, 7d, and 7e). The evident phenotype was observed in *tkr-1 (ok2886)* animals. A Fig. 7b showed significantly low SI values of *tkr-1* KO compared to WT animals in ethanol seeking assay against the Cu ^2+^ barrier in overall concentrations. Remarkably, *tkr-1* KO animals did not cross even low concentration of Cu^2+^ barrier after ethanol pretreatment like *seb-3* KO animals. Interestingly, the KO animals of *tkr-2*, predicted as a NK3R orthologue (^18^), also showed slightly reduced development of aversion-resistant ethanol seeking induced by ethanol pretreatment whereas *tkr-3* KO animals displayed ethanol preference and aversion-resistant ethanol seeking as much as wild type (Fig. 7d, and 7e). Our results, impaired development of aversion-resistant ethanol seeking behavior in the absence of TKR-1, was evident and reflect the importance of the involvement of TKR-1 function in the progress of compulsive ethanol seeking. Combined, these data suggest that SEB-3 transcriptionally regulates the TKR-1 to enhance and progress the compulsive ethanol seeking behavior in ethanol experienced animals.

**Fig. 7.**
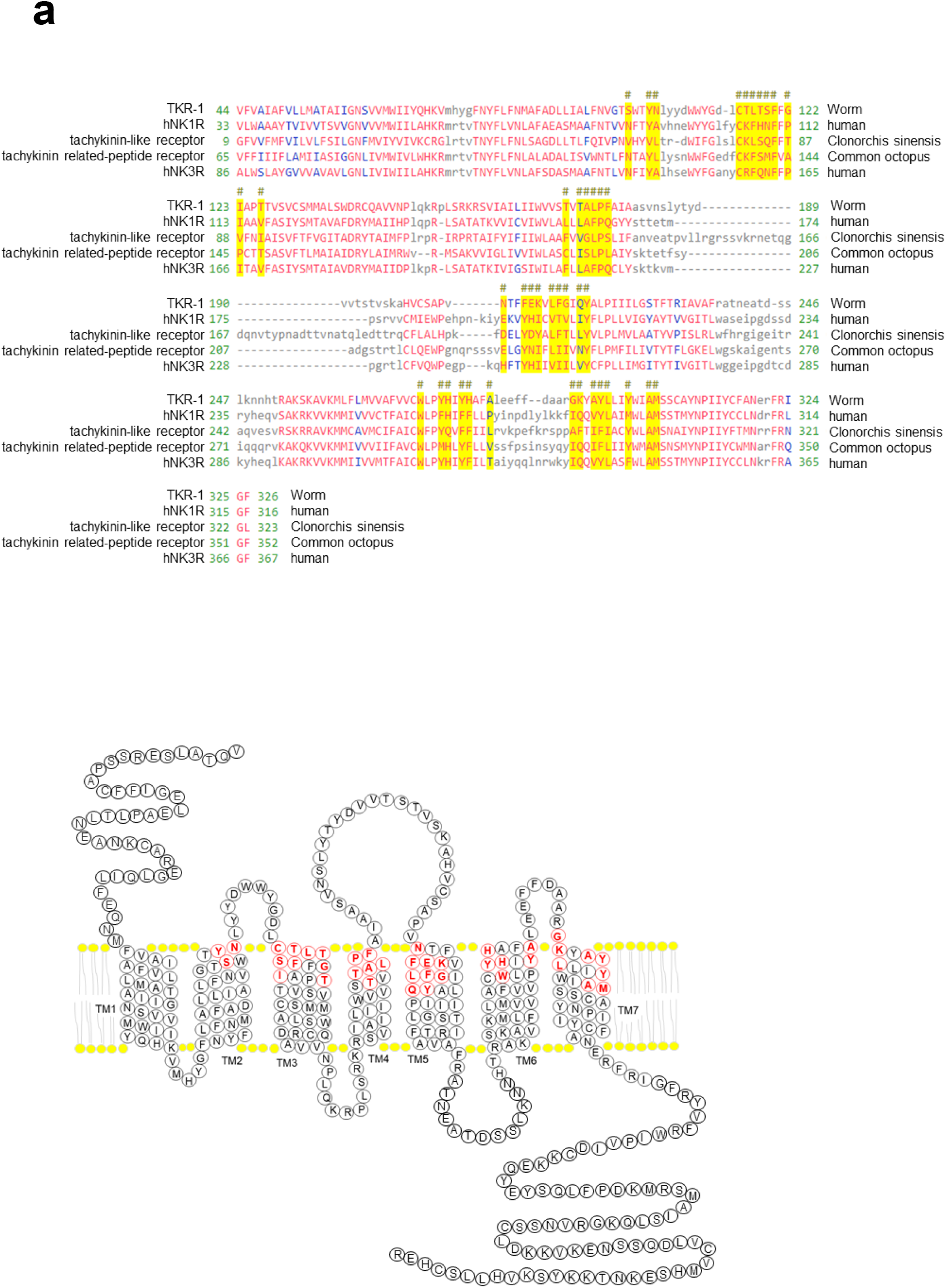
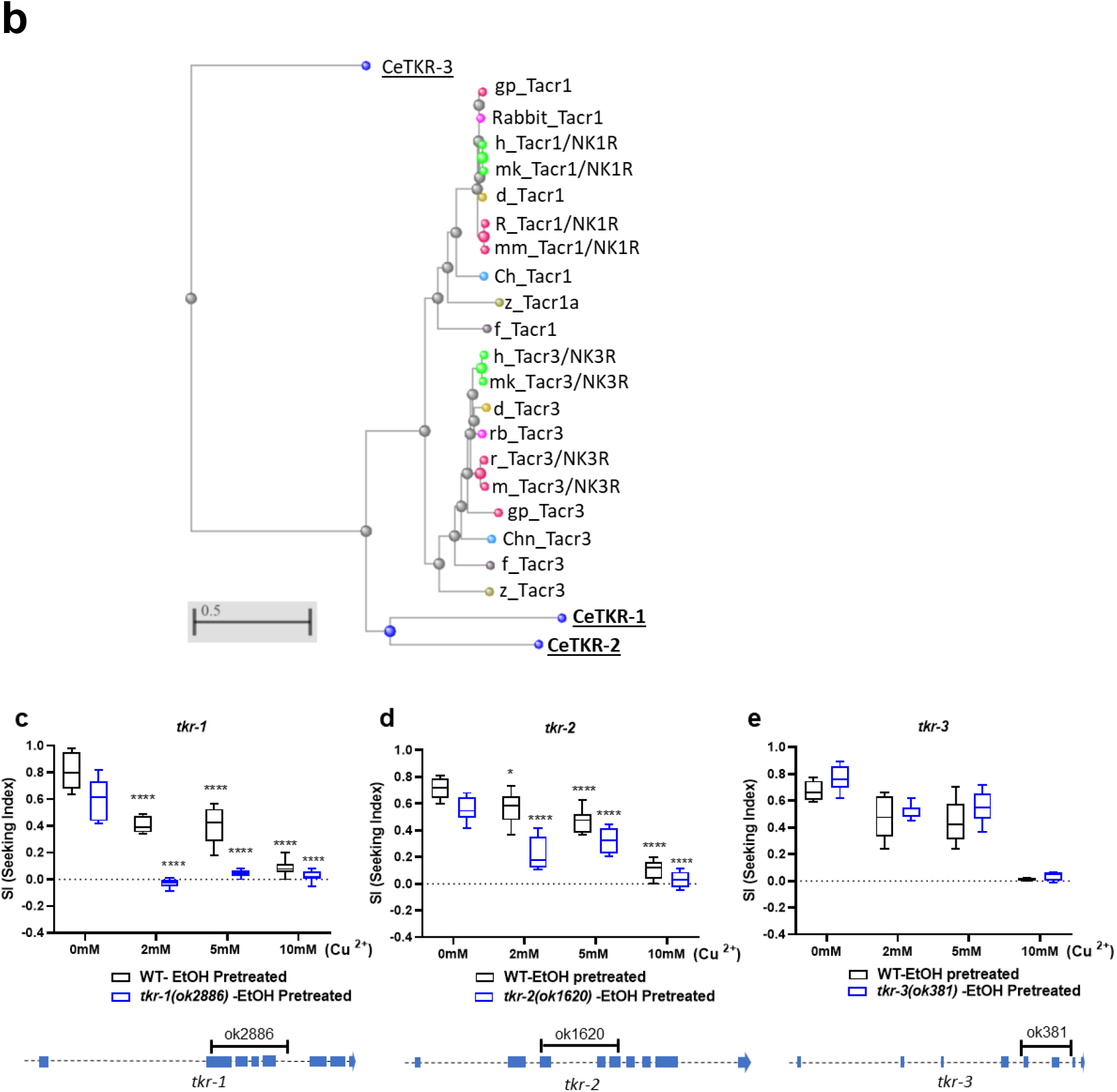
Neurokinin receptors in *C. elegans* (a) Sequence alignment of TKR-1 (NP_499064.2), humanTACR1/NK1R (2KS9_A), humanTACR3/NK3R (P29371), and tachykinin-like receptors of liver fluke (GAA51416) and octopus (BAD93354.1) revealed the consensus sequences of putative ligand binding pocket of TKR-1 (Yellow highlights). Red indicates highly conserved amino acids and blue indicates lower conservation. The diagram of full-length TKR-1 topology represents transmembrane helix domains and the consensus sequences of putative peptide ligand binding pocket (Red amino acids). (b) Phylogenetic analysis of the Tachykinin/Neurokinin Receptor family (minimum evolution method). (c-e) Aversion-resistant ethanol seeking assay in *tkr-1(ok2886)* (c), *tkr-2(ok1620)* (d), and *tkr-3(ok381)* (e). A two-way ANOVA comparison shows the impaired development in *tkr-1(ok2886)* [F_Genotype_(1, 58)=106.6, p<0.0001; F_Concentration_(3, 58)=140.7, p<0.0001; F_Genotype x Concentration_(3, 58)=9.495, p<0.0001]. Significant post hoc differences (Dunnett’s test) between no barrier versus 2mM, 5mM, or 10mM in each genotype (p<0.0001, ****); and *tkr-2(1620)* [F_Genotype_(1, 16)=38.37, p<0.0001; F_Concentration_(3, 34)=76.82, p<0.0001; F_Genotype x Concentration_(3, 34)=6.030, p=0.0021]. Significant post hoc differences (Dunnett’s test) between no barrier versus 2mM, 5mM, or 10mM in each genotype (p<0.05, *; p<0.0001, ****).

## Discussion

Compulsivity is defined as repetitive attempts despite facing adverse consequences and compel individuals to perform to be relieved from stress or anxiety (^68^). The repetitive or obsessive aspects and compulsive drinking scale has been adapted and help to evaluate the severity of AUD in human genetic studies for reliable assessing alcohol craving (^6^, ^26^, ^27^), that has been described as a compelling urge to intake alcohol and considered crucial for the maintenance of AUD (^5^). The compulsive alcohol seeking is characterized by an imbalance between superior drive to alcohol and disruption in control of alcohol use (^12^, ^13^). To model the development of compulsive engagement of alcohol seeking in *C. elegans*, we showed following: i) enhanced preference of ethanol, ii) repetitive attempts to seek, and iii) enhanced aversion-resistant ethanol seeking.

Despite baseline aversive response to ethanol in acute exposure, *C. elegans* exhibit state-dependent ethanol seeking behavior, which is a significantly potentiated preference for ethanol after the prolonged experience of ethanol (^21^). In addition, *C. elegans* also recapitulates the behavioral traits of alcohol-dependent animals, known as studies in mammals. Ethanol pretreated worms showed distinct orientation while seeking for ethanol and a pronounced tendency to stay in the limited region where ethanol is, even though their locomotion and exploratory behavior were not damaged by the pretreatment of ethanol (Fig. 1 and 2). The increase in exploratory behavior during ethanol withdrawal is consistent with the evident demonstration of ethanol-withdrawal symptom in *C. elegans*, as seen in our previous study (^22^), and nicotine, classified as a psychostimulant like ethanol, can also be compared to promoting withdrawal-induced motor stimulation in *C. elegans* (^69^). The locomotion trajectories are depending on the transition between distinct motor states, which is the modulation of random exploration behavior (^70^, ^71^, ^72^, ^73^). Since the long-term behavioral state of exploration in *C. elegans* is modulated by neuropeptide signals including *seb-3* (^74^, ^75^, ^22^), the change in exploration behavior of ethanol pretreated animals can be considered in association with the dependence modulation by neuropeptides signaling.

Ethanol pretreated animals also represented repetitive attempts and endurance to cross the chemical barrier to move to the ethanol area and consequently, crossed the barrier (Fig. 3). Chemotaxis behaviors are regulated primarily by the chemosensory neurons and modulated by integration of signaling with interneurons (^76^, ^77^). Recent studies on multisensory integration of the worms have provided sophisticated experimental manipulation with complex behavioral decision paradigms *in vivo* animals where information from distinct sensory networks is concurrently processed to demonstrate the comprehensible representation of the environment (^78^, ^79^, ^80^). We introduced nociceptive stimuli, of which perception and mediation are well defined (^38^, ^81^, ^82^, ^83^), as an obstacle to interfere with the animals’ ethanol seeking behavior. Since ethanol pretreatment does not change the worm’s sensitivity to the aversive stimulus (Fig.3b), ethanol seeking against the aversive chemical barrier promise as an endophenotype for compulsive ethanol seeking. Together with persistent drug seeking despite aversive consequences, aversion-resistant alcohol intake is increasingly recognized as a pivotal characteristic of addiction and is being investigated as a model of compulsive behavior (^84^, ^85^, ^86^, ^87^). The progress of compulsivity is hypothesized to be imbalanced between enhanced ethanol seeking and loss of controlling avoidance program. *C. elegnas* detect the aversive stimuli in the environment and avoid noxious chemicals to cope with unfavorable situation. After repeated trial, crossing over aversive stimuli for ethanol seeking is depending on the concentration of aversive barrier suggesting the strength of an animal’s innate state that modulate ethanol preference results in compulsive seeking. We introduced various nociceptive chemicals, of which avoidance is mediated by polymodal sensory neurons for nociception (^39^, ^38^), to rule out the possibility that overcoming aversive barrier for ethanol seeking was specific to certain aversive stimuli. For example, in those cases where only the receptors for the copper sensation are downregulated during pretreatment instead of modulation of the overall circuit of attraction or avoidance program depending on the state. Cu^2+^ and denatonium benzoate effectively interfere with ethanol seeking behavior in concentration dependent manner. Quinine also works in a similar way, but only a low concentration was applicable due to the solubility in ethanol as a solvent (data not shown). Together with increased ethanol preference, we demonstrate the ethanol dependent behaviors of *C. elegans* that parallels compulsive alcohol seeking behavior in mammals.

Neuromodulation via neuropeptides signaling has been shown to have a pivotal roles in the development of AUD from worms to human (^20^, ^22^, ^88^, ^89^, ^90^, ^91^, ^33^). We demonstrate that SEB-3, a CRF (corticotropin-releasing factor) receptor-like GPCR in *C. elegans*, positively regulates ethanol preference and also has a role in the progress of compulsive ethanol seeking over adverse stimuli after prolonged preexposure to ethanol. Neuropeptidergic signaling of CRF has been studied as a key element that lead to AUD and comorbidities in a way to compensate for the distress associated with substance withdrawal and discontinuation. Neuropeptides signaling provides powerful mechanisms for rapid physiological adjustment to kaleidoscopic environmental changes and maintain systemic function by precise integration of signal flow. A maladaptation in multisensory integration has been reported in individuals diagnosed with compulsive behaviors (^92^, ^93^). Previously, we have shown insight into how SEB-3 facilitates the progress of compulsive behavior by demonstrating that SEB-3 enhances motivational state and leads animals to engage obsessively in sexual drive, superseding the avoidance program of locomotion under negative stimuli (^94^). The animal has to consider its physiological statuses such as hunger and sexual drive and must monitor its environment to determine whether stressful conditions warrant the expression of their innate urges. The CRF (corticotropin-releasing factor) system plays a pivotal role in mediating stress responses in the brain from amphibians to primates (^95^, ^96^, ^97^, ^98^). A CRF (corticotropin-releasing factor) receptor has been implicated in the pathophysiology of compulsive behavior such as anxiety and AUD (^99^; ^100^; ^101^). Stress pathways have been studied as key fundamentals of neural systems that drive alcohol and drug dependence. The brain stress system is activated and sensitized during repeated withdrawal and lead to a negative emotional state that promotes dependence (^102^). Indeed, the *seb-3(eg696)* gf animals, originally isolated as a genetic variant showing enhanced arousal and represented altered susceptibility to AUD such as enhanced acute functional tolerance to ethanol and withdrawal behavior (^22^).

Like CRF, Tachykinin /Neurokinin is one of the strongest conserved neuropeptide systems across bilaterians, of which both peptide and receptor orthologues are represented in *C*.*elegans* (^58^). The *tkr-1, tkr-2*, and *tkr-3* have been categorized into tachykinin/ neurokinin receptor-like group (^103^, ^104^, ^105^) and RNAi functional screens revealed *tkr-1* affects fat metabolism and deposition (^106^). We report *tkr-1* and *tkr-2*, close to NK1R and NK3R, have a prominent role in the progress of compulsive ethanol seeking. NK1R preferentially mediates the signal of Substance P (SP), belongs to the tachykinin/neurokinin family of neuropeptides (^107^, ^108^). SP and NK1R are widespread in the nervous system in mammals. NK1R has been studied in correlation with the stress response related to anxiety and depression (^62^, ^63^, ^64^). Recently after the failure of CRFR1 antagonist in Clinical trials (^109^, ^110^), the increasing evidence of neurokinin receptors related to alcohol and drug dependence has been more highlighted. An activation of both NK1R and NK3R facilitate dopamine release in the NAc (^111^) and NK1R antagonist was reported to reduce cocaine-induced DA release (^112^). Antagonism of NK1R and KO mice reduces alcohol consumption (^113^) and Antagonism of NK1R decreases alcohol self-administration in alcohol-preferring rats (^114^). Furthermore, adeno-associated virus-mediated overexpression of NK1R in the central amygdala increased alcohol self-administration (^115^), which is consistent with our finding. A Family-based association study in human genetics have also reported NK3R is associated with alcohol and cocaine dependence (^116^).

We report that SEB-3 facilitates animal drive to seek ethanol over noxious stimuli representing enhanced compulsive seeking. Our functional genomics study revealed that TKR-1 is upregulated in *seb-3* gf strain, which is defined as compulsive ethanol seeking animals, and functions in the development of compulsive ethanol seeking. Interestingly, *sodh-1*, which showed a significant alcohol intoxication phenotype in the orthogonal test for functional validation of ADH (alcohol dehydrogenase) as a human GWAS candidate related to heavy alcohol consumption (^25^), is also upregulated in *seb-3* gf animals (Sup.1). These results also demonstrate the potential of our investigation as a scalable model to accelerate the functional validation of alcohol dependence associated with networks of epistatic interactions. Despite strong evidence of the crucial role of the CRF system in the development of alcohol dependence, there are concerns about the importance of the CRF system for drug target based on recent failures of CRF1 antagonist clinical trials for alcohol dependence. Here, the conservation of ethanol phenotypes across species suggests that a gradual increase of dependence-induced compulsivity is progressed by the interplay between CRF and Neurokinin signaling. Further investigation of interactions will provide significance to understand the neurobiological mechanism to progress AUD for advanced clinical benefits.

## Acknowledgments

This work was supported by College of Medicine, University of Tennessee Health Science Center (UTHSC). We thank the *C. elegans* Genetics Center (CGC) for providing strains, which is funded by NIH Office of Research Infrastructure Programs (P40 OD010440). Authors declare no conflict of interests.

