## Supplemental.1 for "Neuropeptidergic regulation of Compulsive Ethanol Seeking in *C. elegans*"

**List of genes associated with GO terms in Fig. 6b**

| GO_term | gene |
|---|---|
| defense response GO:0006952 | WBGene00003766 |
| defense response GO:0006952 | WBGene00000556 |
| defense response GO:0006952 | WBGene00044719 |
| defense response GO:0006952 | WBGene00015759 |
| defense response GO:0006952 | WBGene00003767 |
| defense response GO:0006952 | WBGene00009226 |
| defense response GO:0006952 | WBGene00018731 |
| defense response GO:0006952 | WBGene00019298 |
| defense response GO:0006952 | WBGene00018384 |
| defense response GO:0006952 | WBGene00004986 |
| defense response GO:0006952 | WBGene00010790 |
| defense response GO:0006952 | WBGene00012783 |
| defense response GO:0006952 | WBGene00003768 |
| defense response GO:0006952 | WBGene00017364 |
| defense response GO:0006952 | WBGene00009429 |
| defense response GO:0006952 | WBGene00000169 |
| defense response GO:0006952 | WBGene00003765 |
| response to biotic stimulus GO:0009607 | WBGene00003766 |
| response to biotic stimulus GO:0009607 | WBGene00000556 |
| response to biotic stimulus GO:0009607 | WBGene00044719 |
| response to biotic stimulus GO:0009607 | WBGene00015759 |
| response to biotic stimulus GO:0009607 | WBGene00003767 |
| response to biotic stimulus GO:0009607 | WBGene00009226 |
| response to biotic stimulus GO:0009607 | WBGene00018731 |
| response to biotic stimulus GO:0009607 | WBGene00019298 |
| response to biotic stimulus GO:0009607 | WBGene00018384 |
| response to biotic stimulus GO:0009607 | WBGene00004986 |
| response to biotic stimulus GO:0009607 | WBGene00010790 |
| response to biotic stimulus GO:0009607 | WBGene00012783 |
| response to biotic stimulus GO:0009607 | WBGene00003768 |
| response to biotic stimulus GO:0009607 | WBGene00017364 |
| response to biotic stimulus GO:0009607 | WBGene00009429 |
| response to biotic stimulus GO:0009607 | WBGene00000169 |
| response to biotic stimulus GO:0009607 | WBGene00003765 |
| neuropeptide signaling pathway GO:0007218 | WBGene00003766 |
| neuropeptide signaling pathway GO:0007218 | WBGene00015046 |
| neuropeptide signaling pathway GO:0007218 | WBGene00016984 |
| neuropeptide signaling pathway GO:0007218 | WBGene00003767 |
| neuropeptide signaling pathway GO:0007218 | WBGene00003768 |
| neuropeptide signaling pathway GO:0007218 | WBGene00006576 |
| neuropeptide signaling pathway GO:0007218 | WBGene00003765 |

| | |
|---|---|
| organic acid metabolic process GO:0006082 | WBGene00007836 |
| organic acid metabolic process GO:0006082 | WBGene00016768 |
| organic acid metabolic process GO:0006082 | WBGene00012608 |
| organic acid metabolic process GO:0006082 | WBGene00019492 |
| organic acid metabolic process GO:0006082 | WBGene00017565 |
| organic acid metabolic process GO:0006082 | WBGene00009334 |
| organic acid metabolic process GO:0006082 | WBGene00014182 |
| organic acid metabolic process GO:0006082 | WBGene00004258 |
| organic acid metabolic process GO:0006082 | WBGene00016630 |
| organic acid metabolic process GO:0006082 | WBGene00001564 |
| organic acid metabolic process GO:0006082 | WBGene00010759 |
| organic acid metabolic process GO:0006082 | WBGene00015044 |
| organic acid metabolic process GO:0006082 | WBGene00018286 |
| tetrapyrrole binding GO:0046906 | WBGene00000831 |
| tetrapyrrole binding GO:0046906 | WBGene00008519 |
| tetrapyrrole binding GO:0046906 | WBGene00016768 |
| tetrapyrrole binding GO:0046906 | WBGene00009226 |
| tetrapyrrole binding GO:0046906 | WBGene00011287 |
| tetrapyrrole binding GO:0046906 | WBGene00015044 |
| iron ion binding GO:0005506 | WBGene00008519 |
| iron ion binding GO:0005506 | WBGene00016768 |
| iron ion binding GO:0005506 | WBGene00001523 |
| iron ion binding GO:0005506 | WBGene00009226 |
| iron ion binding GO:0005506 | WBGene00015044 |
| dendritic tree GO:0097447 | WBGene00006412 |
| dendritic tree GO:0097447 | WBGene00000100 |
| dendritic tree GO:0097447 | WBGene00003856 |
| dendritic tree GO:0097447 | WBGene00001121 |
| dendritic tree GO:0097447 | WBGene00015350 |
| cation binding GO:0043169 | WBGene00000245 |
| cation binding GO:0043169 | WBGene00003528 |
| cation binding GO:0043169 | WBGene00003634 |
| cation binding GO:0043169 | WBGene00000831 |
| cation binding GO:0043169 | WBGene00008519 |
| cation binding GO:0043169 | WBGene00016768 |
| cation binding GO:0043169 | WBGene00012608 |
| cation binding GO:0043169 | WBGene00003689 |
| cation binding GO:0043169 | WBGene00001523 |
| cation binding GO:0043169 | WBGene00016903 |
| cation binding GO:0043169 | WBGene00009226 |
| cation binding GO:0043169 | WBGene00010681 |
| cation binding GO:0043169 | WBGene00003618 |
| cation binding GO:0043169 | WBGene00000100 |

| | |
|---|---|
| cation binding GO:0043169 | WBGene00018384 |
| cation binding GO:0043169 | WBGene00015332 |
| cation binding GO:0043169 | WBGene00003693 |
| cation binding GO:0043169 | WBGene00004258 |
| cation binding GO:0043169 | WBGene00010790 |
| cation binding GO:0043169 | WBGene00009503 |
| cation binding GO:0043169 | WBGene00001121 |
| cation binding GO:0043169 | WBGene00001184 |
| cation binding GO:0043169 | WBGene00015044 |
| cation binding GO:0043169 | WBGene00003695 |
| cation binding GO:0043169 | WBGene00011003 |
| cation binding GO:0043169 | WBGene00003544 |
| cation binding GO:0043169 | WBGene00017340 |
| cation binding GO:0043169 | WBGene00000288 |
| passive transmembrane transporter activity GO:0022803 | WBGene00001373 |
| passive transmembrane transporter activity GO:0022803 | WBGene00006784 |
| passive transmembrane transporter activity GO:0022803 | WBGene00006614 |
| passive transmembrane transporter activity GO:0022803 | WBGene00004986 |
| passive transmembrane transporter activity GO:0022803 | WBGene00001171 |
| passive transmembrane transporter activity GO:0022803 | WBGene00001591 |
| passive transmembrane transporter activity GO:0022803 | WBGene00000176 |
| passive transmembrane transporter activity GO:0022803 | WBGene00000169 |
| passive transmembrane transporter activity GO:0022803 | WBGene00007808 |
| transmembrane transport GO:0055085 | WBGene00004878 |
| transmembrane transport GO:0055085 | WBGene00001373 |
| transmembrane transport GO:0055085 | WBGene00013499 |
| transmembrane transport GO:0055085 | WBGene00020700 |
| transmembrane transport GO:0055085 | WBGene00006784 |
| transmembrane transport GO:0055085 | WBGene00018779 |
| transmembrane transport GO:0055085 | WBGene00006614 |
| transmembrane transport GO:0055085 | WBGene00012259 |
| transmembrane transport GO:0055085 | WBGene00004986 |
| transmembrane transport GO:0055085 | WBGene00017663 |
| transmembrane transport GO:0055085 | WBGene00001171 |
| transmembrane transport GO:0055085 | WBGene00019300 |
| transmembrane transport GO:0055085 | WBGene00001591 |
| transmembrane transport GO:0055085 | WBGene00000176 |
| transmembrane transport GO:0055085 | WBGene00000169 |
| transmembrane transport GO:0055085 | WBGene00016628 |
| transmembrane transport GO:0055085 | WBGene00007808 |
| response to endogenous stimulus GO:0009719 | WBGene00020649 |
| response to endogenous stimulus GO:0009719 | WBGene00003618 |
| response to endogenous stimulus GO:0009719 | WBGene00001184 |

| | | |
|---|---|---|
| response to endogenous stimulus | GO:0009719 | WBGene00002163 |
| response to endogenous stimulus | GO:0009719 | WBGene00003695 |
