## Supplemental. 2 for "Neuropeptidergic regulation of Compulsive Ethanol Seeking in *C. elegans*"

**List of genes associated with GO terms in Fig. 6c**

| GO_term | gene |
|---|---|
| immune system process GO:0002376 | WBGene00019213 |
| immune system process GO:0002376 | WBGene00003091 |
| immune system process GO:0002376 | WBGene00007440 |
| immune system process GO:0002376 | WBGene00010470 |
| immune system process GO:0002376 | WBGene00012462 |
| immune system process GO:0002376 | WBGene00018823 |
| immune system process GO:0002376 | WBGene00009710 |
| immune system process GO:0002376 | WBGene00008486 |
| immune system process GO:0002376 | WBGene00008199 |
| immune system process GO:0002376 | WBGene00009431 |
| immune system process GO:0002376 | WBGene00003995 |
| immune system process GO:0002376 | WBGene00044457 |
| immune system process GO:0002376 | WBGene00015933 |
| immune system process GO:0002376 | WBGene00015362 |
| immune system process GO:0002376 | WBGene00003090 |
| immune system process GO:0002376 | WBGene00007506 |
| immune system process GO:0002376 | WBGene00012101 |
| immune system process GO:0002376 | WBGene00219216 |
| immune system process GO:0002376 | WBGene00021518 |
| immune system process GO:0002376 | WBGene00003096 |
| immune system process GO:0002376 | WBGene00009971 |
| immune system process GO:0002376 | WBGene00015216 |
| immune system process GO:0002376 | WBGene00008900 |
| immune system process GO:0002376 | WBGene00008631 |
| immune system process GO:0002376 | WBGene00010125 |
| immune system process GO:0002376 | WBGene00000522 |
| immune system process GO:0002376 | WBGene00020455 |
| immune system process GO:0002376 | WBGene00007097 |
| immune system process GO:0002376 | WBGene00016788 |
| immune system process GO:0002376 | WBGene00010745 |
| immune system process GO:0002376 | WBGene00015455 |
| immune system process GO:0002376 | WBGene00010659 |
| immune system process GO:0002376 | WBGene00008988 |
| immune system process GO:0002376 | WBGene00022848 |
| immune system process GO:0002376 | WBGene00001772 |
| immune system process GO:0002376 | WBGene00009914 |
| immune system process GO:0002376 | WBGene00003094 |
| immune system process GO:0002376 | WBGene00009520 |
| immune system process GO:0002376 | WBGene00021519 |
| immune system process GO:0002376 | WBGene00007153 |
| immune system process GO:0002376 | WBGene00012947 |
| immune system process GO:0002376 | WBGene00015716 |
| immune system process GO:0002376 | WBGene00015259 |
| immune system process GO:0002376 | WBGene00011979 |
| immune system process GO:0002376 | WBGene00015932 |

| | |
|---|---|
| immune system process GO:0002376 | WBGene00019660 |
| immune system process GO:0002376 | WBGene00008726 |
| immune system process GO:0002376 | WBGene00019564 |
| immune system process GO:0002376 | WBGene00010658 |
| immune system process GO:0002376 | WBGene00001761 |
| immune system process GO:0002376 | WBGene00000841 |
| immune system process GO:0002376 | WBGene00016912 |
| immune system process GO:0002376 | WBGene00016425 |
| immune system process GO:0002376 | WBGene00018643 |
| immune system process GO:0002376 | WBGene00020579 |
| immune system process GO:0002376 | WBGene00007717 |
| immune system process GO:0002376 | WBGene00020799 |
| immune system process GO:0002376 | WBGene00009526 |
| immune system process GO:0002376 | WBGene00008577 |
| immune system process GO:0002376 | WBGene00017726 |
| immune system process GO:0002376 | WBGene00020760 |
| immune system process GO:0002376 | WBGene00002274 |
| immune system process GO:0002376 | WBGene00001749 |
| response to biotic stimulus GO:0009607 | WBGene00019213 |
| response to biotic stimulus GO:0009607 | WBGene00003091 |
| response to biotic stimulus GO:0009607 | WBGene00008199 |
| response to biotic stimulus GO:0009607 | WBGene00009431 |
| response to biotic stimulus GO:0009607 | WBGene00003995 |
| response to biotic stimulus GO:0009607 | WBGene00015933 |
| response to biotic stimulus GO:0009607 | WBGene00003090 |
| response to biotic stimulus GO:0009607 | WBGene00003096 |
| response to biotic stimulus GO:0009607 | WBGene00009971 |
| response to biotic stimulus GO:0009607 | WBGene00015052 |
| response to biotic stimulus GO:0009607 | WBGene00003093 |
| response to biotic stimulus GO:0009607 | WBGene00010125 |
| response to biotic stimulus GO:0009607 | WBGene00000522 |
| response to biotic stimulus GO:0009607 | WBGene00008988 |
| response to biotic stimulus GO:0009607 | WBGene00003094 |
| response to biotic stimulus GO:0009607 | WBGene00015932 |
| response to biotic stimulus GO:0009607 | WBGene00000841 |
| response to biotic stimulus GO:0009607 | WBGene00018643 |
| response to biotic stimulus GO:0009607 | WBGene00020579 |
| response to biotic stimulus GO:0009607 | WBGene00012211 |
| response to biotic stimulus GO:0009607 | WBGene00001749 |
| **membrane GO:0016020** | WBGene00019213 |
| **membrane GO:0016020** | WBGene00016481 |
| **membrane GO:0016020** | WBGene00021743 |
| **membrane GO:0016020** | WBGene00009644 |
| **membrane GO:0016020** | WBGene00008647 |
| **membrane GO:0016020** | WBGene00019200 |
| **membrane GO:0016020** | WBGene00018823 |
| **membrane GO:0016020** | WBGene00009710 |

| | |
|---|---|
| **membrane GO:0016020** | WBGene00000693 |
| **membrane GO:0016020** | WBGene00017069 |
| **membrane GO:0016020** | WBGene00008486 |
| **membrane GO:0016020** | WBGene00019146 |
| **membrane GO:0016020** | WBGene00009431 |
| **membrane GO:0016020** | WBGene00003995 |
| **membrane GO:0016020** | WBGene00044457 |
| **membrane GO:0016020** | WBGene00194926 |
| **membrane GO:0016020** | WBGene00013602 |
| **membrane GO:0016020** | WBGene00013586 |
| **membrane GO:0016020** | WBGene00008850 |
| **membrane GO:0016020** | WBGene00020120 |
| **membrane GO:0016020** | WBGene00015933 |
| **membrane GO:0016020** | WBGene00000133 |
| **membrane GO:0016020** | WBGene00006947 |
| **membrane GO:0016020** | WBGene00021339 |
| **membrane GO:0016020** | WBGene00004002 |
| **membrane GO:0016020** | WBGene00011124 |
| **membrane GO:0016020** | WBGene00021948 |
| **membrane GO:0016020** | WBGene00012293 |
| **membrane GO:0016020** | WBGene00015965 |
| **membrane GO:0016020** | WBGene00021518 |
| **membrane GO:0016020** | WBGene00013073 |
| **membrane GO:0016020** | WBGene00194730 |
| **membrane GO:0016020** | WBGene00015152 |
| **membrane GO:0016020** | WBGene00011668 |
| **membrane GO:0016020** | WBGene00010249 |
| **membrane GO:0016020** | WBGene00017329 |
| **membrane GO:0016020** | WBGene00077525 |
| **membrane GO:0016020** | WBGene00011190 |
| **membrane GO:0016020** | WBGene00017948 |
| **membrane GO:0016020** | WBGene00009971 |
| **membrane GO:0016020** | WBGene00015052 |
| **membrane GO:0016020** | WBGene00016013 |
| **membrane GO:0016020** | WBGene00003573 |
| **membrane GO:0016020** | WBGene00008631 |
| **membrane GO:0016020** | WBGene00005586 |
| **membrane GO:0016020** | WBGene00015879 |
| **membrane GO:0016020** | WBGene00009488 |
| **membrane GO:0016020** | WBGene00022401 |
| **membrane GO:0016020** | WBGene00000522 |
| **membrane GO:0016020** | WBGene00001817 |
| **membrane GO:0016020** | WBGene00015692 |
| **membrane GO:0016020** | WBGene00016909 |
| **membrane GO:0016020** | WBGene00020587 |
| **membrane GO:0016020** | WBGene00015080 |
| **membrane GO:0016020** | WBGene00019550 |

| | |
|---|---|
| **membrane GO:0016020** | WBGene00000675 |
| **membrane GO:0016020** | WBGene00004353 |
| **membrane GO:0016020** | WBGene00002090 |
| **membrane GO:0016020** | WBGene00008084 |
| **membrane GO:0016020** | WBGene00010157 |
| **membrane GO:0016020** | WBGene00001536 |
| **membrane GO:0016020** | WBGene00007097 |
| **membrane GO:0016020** | WBGene00021448 |
| **membrane GO:0016020** | WBGene00019234 |
| **membrane GO:0016020** | WBGene00015455 |
| **membrane GO:0016020** | WBGene00021034 |
| **membrane GO:0016020** | WBGene00008988 |
| **membrane GO:0016020** | WBGene00194747 |
| **membrane GO:0016020** | WBGene00016797 |
| **membrane GO:0016020** | WBGene00000716 |
| **membrane GO:0016020** | WBGene00018335 |
| **membrane GO:0016020** | WBGene00015449 |
| **membrane GO:0016020** | WBGene00021797 |
| **membrane GO:0016020** | WBGene00009914 |
| **membrane GO:0016020** | WBGene00015386 |
| **membrane GO:0016020** | WBGene00005188 |
| **membrane GO:0016020** | WBGene00004003 |
| **membrane GO:0016020** | WBGene00021519 |
| **membrane GO:0016020** | WBGene00013728 |
| **membrane GO:0016020** | WBGene00020182 |
| **membrane GO:0016020** | WBGene00000696 |
| **membrane GO:0016020** | WBGene00002268 |
| **membrane GO:0016020** | WBGene00005653 |
| **membrane GO:0016020** | WBGene00012947 |
| **membrane GO:0016020** | WBGene00008296 |
| **membrane GO:0016020** | WBGene00017168 |
| **membrane GO:0016020** | WBGene00015259 |
| **membrane GO:0016020** | WBGene00013874 |
| **membrane GO:0016020** | WBGene00022736 |
| **membrane GO:0016020** | WBGene00009859 |
| **membrane GO:0016020** | WBGene00015932 |
| **membrane GO:0016020** | WBGene00016285 |
| **membrane GO:0016020** | WBGene00012552 |
| **membrane GO:0016020** | WBGene00019660 |
| **membrane GO:0016020** | WBGene00005555 |
| **membrane GO:0016020** | WBGene00045416 |
| **membrane GO:0016020** | WBGene00010296 |
| **membrane GO:0016020** | WBGene00013540 |
| **membrane GO:0016020** | WBGene00016425 |
| **membrane GO:0016020** | WBGene00010998 |
| **membrane GO:0016020** | WBGene00005269 |
| **membrane GO:0016020** | WBGene00013484 |

| | |
|---|---|
| **membrane GO:0016020** | WBGene00020579 |
| **membrane GO:0016020** | WBGene00010206 |
| **membrane GO:0016020** | WBGene00007717 |
| **membrane GO:0016020** | WBGene00010749 |
| **membrane GO:0016020** | WBGene00020799 |
| **membrane GO:0016020** | WBGene00021153 |
| **membrane GO:0016020** | WBGene00000757 |
| **membrane GO:0016020** | WBGene00016063 |
| **membrane GO:0016020** | WBGene00012163 |
| **membrane GO:0016020** | WBGene00019121 |
| **membrane GO:0016020** | WBGene00006272 |
| **membrane GO:0016020** | WBGene00020760 |
| **membrane GO:0016020** | WBGene00012222 |
| **membrane GO:0016020** | WBGene00007829 |
| **membrane GO:0016020** | WBGene00012323 |
| **membrane GO:0016020** | WBGene00004979 |
| **membrane GO:0016020** | WBGene00009713 |
| **membrane GO:0016020** | WBGene00022411 |
| **membrane GO:0016020** | WBGene00010362 |
| **intrinsic component of membrane GO:0031224** | WBGene00019213 |
| **intrinsic component of membrane GO:0031224** | WBGene00016481 |
| **intrinsic component of membrane GO:0031224** | WBGene00021743 |
| **intrinsic component of membrane GO:0031224** | WBGene00009644 |
| **intrinsic component of membrane GO:0031224** | WBGene00008647 |
| **intrinsic component of membrane GO:0031224** | WBGene00019200 |
| **intrinsic component of membrane GO:0031224** | WBGene00009710 |
| **intrinsic component of membrane GO:0031224** | WBGene00000693 |
| **intrinsic component of membrane GO:0031224** | WBGene00017069 |
| **intrinsic component of membrane GO:0031224** | WBGene00008486 |
| **intrinsic component of membrane GO:0031224** | WBGene00019146 |
| **intrinsic component of membrane GO:0031224** | WBGene00003995 |
| **intrinsic component of membrane GO:0031224** | WBGene00044457 |
| **intrinsic component of membrane GO:0031224** | WBGene00194926 |
| **intrinsic component of membrane GO:0031224** | WBGene00013602 |
| **intrinsic component of membrane GO:0031224** | WBGene00013586 |
| **intrinsic component of membrane GO:0031224** | WBGene00008850 |
| **intrinsic component of membrane GO:0031224** | WBGene00020120 |
| **intrinsic component of membrane GO:0031224** | WBGene00000133 |
| **intrinsic component of membrane GO:0031224** | WBGene00021339 |
| **intrinsic component of membrane GO:0031224** | WBGene00004002 |
| **intrinsic component of membrane GO:0031224** | WBGene00011124 |
| **intrinsic component of membrane GO:0031224** | WBGene00021948 |
| **intrinsic component of membrane GO:0031224** | WBGene00012293 |
| **intrinsic component of membrane GO:0031224** | WBGene00015965 |
| **intrinsic component of membrane GO:0031224** | WBGene00013073 |
| **intrinsic component of membrane GO:0031224** | WBGene00194730 |
| **intrinsic component of membrane GO:0031224** | WBGene00015152 |

| | |
|---|---|
| **intrinsic component of membrane GO:0031224** | WBGene00011668 |
| **intrinsic component of membrane GO:0031224** | WBGene00010249 |
| **intrinsic component of membrane GO:0031224** | WBGene00017329 |
| **intrinsic component of membrane GO:0031224** | WBGene00077525 |
| **intrinsic component of membrane GO:0031224** | WBGene00011190 |
| **intrinsic component of membrane GO:0031224** | WBGene00017948 |
| **intrinsic component of membrane GO:0031224** | WBGene00015052 |
| **intrinsic component of membrane GO:0031224** | WBGene00016013 |
| **intrinsic component of membrane GO:0031224** | WBGene00003573 |
| **intrinsic component of membrane GO:0031224** | WBGene00008631 |
| **intrinsic component of membrane GO:0031224** | WBGene00005586 |
| **intrinsic component of membrane GO:0031224** | WBGene00015879 |
| **intrinsic component of membrane GO:0031224** | WBGene00009488 |
| **intrinsic component of membrane GO:0031224** | WBGene00022401 |
| **intrinsic component of membrane GO:0031224** | WBGene00000522 |
| **intrinsic component of membrane GO:0031224** | WBGene00001817 |
| **intrinsic component of membrane GO:0031224** | WBGene00015692 |
| **intrinsic component of membrane GO:0031224** | WBGene00016909 |
| **intrinsic component of membrane GO:0031224** | WBGene00020587 |
| **intrinsic component of membrane GO:0031224** | WBGene00015080 |
| **intrinsic component of membrane GO:0031224** | WBGene00019550 |
| **intrinsic component of membrane GO:0031224** | WBGene00000675 |
| **intrinsic component of membrane GO:0031224** | WBGene00002090 |
| **intrinsic component of membrane GO:0031224** | WBGene00008084 |
| **intrinsic component of membrane GO:0031224** | WBGene00010157 |
| **intrinsic component of membrane GO:0031224** | WBGene00001536 |
| **intrinsic component of membrane GO:0031224** | WBGene00021448 |
| **intrinsic component of membrane GO:0031224** | WBGene00019234 |
| **intrinsic component of membrane GO:0031224** | WBGene00015455 |
| **intrinsic component of membrane GO:0031224** | WBGene00021034 |
| **intrinsic component of membrane GO:0031224** | WBGene00194747 |
| **intrinsic component of membrane GO:0031224** | WBGene00016797 |
| **intrinsic component of membrane GO:0031224** | WBGene00000716 |
| **intrinsic component of membrane GO:0031224** | WBGene00018335 |
| **intrinsic component of membrane GO:0031224** | WBGene00015449 |
| **intrinsic component of membrane GO:0031224** | WBGene00021797 |
| **intrinsic component of membrane GO:0031224** | WBGene00009914 |
| **intrinsic component of membrane GO:0031224** | WBGene00015386 |
| **intrinsic component of membrane GO:0031224** | WBGene00005188 |
| **intrinsic component of membrane GO:0031224** | WBGene00004003 |
| **intrinsic component of membrane GO:0031224** | WBGene00013728 |
| **intrinsic component of membrane GO:0031224** | WBGene00020182 |
| **intrinsic component of membrane GO:0031224** | WBGene00000696 |
| **intrinsic component of membrane GO:0031224** | WBGene00005653 |
| **intrinsic component of membrane GO:0031224** | WBGene00008296 |
| **intrinsic component of membrane GO:0031224** | WBGene00017168 |
| **intrinsic component of membrane GO:0031224** | WBGene00013874 |

| | |
|---|---|
| **intrinsic component of membrane GO:0031224** | WBGene00022736 |
| **intrinsic component of membrane GO:0031224** | WBGene00009859 |
| **intrinsic component of membrane GO:0031224** | WBGene00016285 |
| **intrinsic component of membrane GO:0031224** | WBGene00012552 |
| **intrinsic component of membrane GO:0031224** | WBGene00005555 |
| **intrinsic component of membrane GO:0031224** | WBGene00045416 |
| **intrinsic component of membrane GO:0031224** | WBGene00010296 |
| **intrinsic component of membrane GO:0031224** | WBGene00013540 |
| **intrinsic component of membrane GO:0031224** | WBGene00010998 |
| **intrinsic component of membrane GO:0031224** | WBGene00005269 |
| **intrinsic component of membrane GO:0031224** | WBGene00013484 |
| **intrinsic component of membrane GO:0031224** | WBGene00020579 |
| **intrinsic component of membrane GO:0031224** | WBGene00010206 |
| **intrinsic component of membrane GO:0031224** | WBGene00007717 |
| **intrinsic component of membrane GO:0031224** | WBGene00010749 |
| **intrinsic component of membrane GO:0031224** | WBGene00020799 |
| **intrinsic component of membrane GO:0031224** | WBGene00021153 |
| **intrinsic component of membrane GO:0031224** | WBGene00000757 |
| **intrinsic component of membrane GO:0031224** | WBGene00016063 |
| **intrinsic component of membrane GO:0031224** | WBGene00012163 |
| **intrinsic component of membrane GO:0031224** | WBGene00006272 |
| **intrinsic component of membrane GO:0031224** | WBGene00020760 |
| **intrinsic component of membrane GO:0031224** | WBGene00012222 |
| **intrinsic component of membrane GO:0031224** | WBGene00007829 |
| **intrinsic component of membrane GO:0031224** | WBGene00012323 |
| **intrinsic component of membrane GO:0031224** | WBGene00004979 |
| **intrinsic component of membrane GO:0031224** | WBGene00009713 |
| **intrinsic component of membrane GO:0031224** | WBGene00022411 |
| **intrinsic component of membrane GO:0031224** | WBGene00010362 |
| protein catabolic process GO:0030163 | WBGene00021743 |
| protein catabolic process GO:0030163 | WBGene00012683 |
| protein catabolic process GO:0030163 | WBGene00020611 |
| protein catabolic process GO:0030163 | WBGene00000784 |
| protein catabolic process GO:0030163 | WBGene00012682 |
| protein catabolic process GO:0030163 | WBGene00015879 |
| protein catabolic process GO:0030163 | WBGene00020609 |
| protein catabolic process GO:0030163 | WBGene00000841 |
| protein catabolic process GO:0030163 | WBGene00019121 |
| protein catabolic process GO:0030163 | WBGene00007605 |
| organic acid metabolic process GO:0006082 | WBGene00010456 |
| organic acid metabolic process GO:0006082 | WBGene00007507 |
| organic acid metabolic process GO:0006082 | WBGene00008435 |
| organic acid metabolic process GO:0006082 | WBGene00013602 |
| organic acid metabolic process GO:0006082 | WBGene00007857 |
| organic acid metabolic process GO:0006082 | WBGene00009048 |
| organic acid metabolic process GO:0006082 | WBGene00001604 |
| organic acid metabolic process GO:0006082 | WBGene00019404 |

| | |
|---|---|
| organic acid metabolic process GO:0006082 | WBGene00003844 |
| organic acid metabolic process GO:0006082 | WBGene00016201 |
| organic acid metabolic process GO:0006082 | WBGene00001158 |
| collagen trimer GO:0005581 | WBGene00000693 |
| collagen trimer GO:0005581 | WBGene00000708 |
| collagen trimer GO:0005581 | WBGene00000675 |
| collagen trimer GO:0005581 | WBGene00000716 |
| collagen trimer GO:0005581 | WBGene00000696 |
| collagen trimer GO:0005581 | WBGene00000757 |
